# Novel *Drosophila cis*-regulatory elements can be uncovered by footprinting transcription factor binding sites in ATAC-seq data

**DOI:** 10.64898/2026.06.22.733832

**Authors:** Christian Mei, Jillian Ness, Katherine Nakai, Zeba Wunderlich

## Abstract

Developmental processes depend on carefully coordinated gene expression. Expression is modulated by the binding of transcription factors (TFs) to cis-regulatory elements (CREs), like enhancers and promoters. Many computational and experimental approaches have been developed to find CREs, particularly enhancers, in the genome, each with strengths and caveats. Given the increasing availability of ATAC-seq data and methods to find TF binding therein, we hypothesized that we could use TF footprinting tools to find clusters of TF binding events within accessible chromatin that may act as CREs. Using *Drosophila* anterior-posterior patterning network as a test bed, we used a digital genomic footprinting tool (DGT), TOBIAS, on previously published early embryo ATAC-seq data to characterize the TF footprint landscape of 16 TFs essential for embryonic patterning. Even in this system, with its extensive enhancer annotation, most footprinted TF binding sites lie outside of known enhancers, with intergenic and intronic regions hosting the highest TF footprint count, albeit at low density. To find potential novel enhancers, we identified high-density TF footprint clusters that are highly conserved and overlap with active enhancer histone mark signals. Five high confidence candidates were selected for reporter assay validation and all five were found to drive spatially patterned expression in the embryo. This study shows that even in a highly characterized system, the analysis of footprinted TF binding sites in ATAC-seq data can uncover new regulatory regions and suggests this approach may be helpful in using existing ATAC-seq data to find novel CREs.

**ARTICLE SUMMARY:** Given the increasing availability of ATAC-seq datasets, workflows to exploit the data to uncover new cis-regulatory elements (CREs), including enhancers, are valuable. Using early anterior-posterior patterning in the *Drosophila* embryo as a test case, we find that previously published transcription factor footprinting tools and ATAC-seq data can be analyzed to yield new candidate CREs. Experimental validation confirms the activity of selected candidate CREs, suggesting that existing data can be analyzed to find novel regulatory elements.

## INTRODUCTION

Development is a complex process that requires careful regulation of gene expression at each step of transcription, translation, and beyond. Control at the first step in gene expression, transcription initiation, is guided by multiple transcription factors (TFs) that bind to specific regions of the genome called *cis-*regulatory elements (CREs), activating or repressing target genes needed for accurate development (Long et al. 2016).

Due to the degenerate binding preferences of eukaryotic TFs (Wunderlich and Mirny 2009), finding all the functional TF binding sites and CREs in the genome has been a long-standing challenge of the field. TF binding preferences can be measured in vitro via protein binding microarrays, SELEX, and other techniques to yield binding motifs (Lambert et al. 2018). Using these motifs, there are numerous computational methods to find clusters of predicted TF binding sites that may be CREs (Hardison and Taylor 2012). However, these methods can yield false positives due to factors that affect in vivo TF binding, like DNA shape, DNA methylation, chromatin accessibility, and histone modifications (Jayaram et al. 2016; Lambert et al. 2018).

Some of these caveats were addressed with the advent of in vivo technologies like chromatin immunoprecipitation followed by sequencing (ChIP-seq) (Johnson et al. 2007), which has been used extensively to map TF binding sites and histone modifications to identify novel CREs (Gasperini et al. 2020). However, ChIP-seq based mapping is limited to one TF or histone modification at a time and is dependent on the availability of suitable antibodies.

With increasing frequency, chromatin accessibility measured by DNase-seq and ATAC-seq has been used to assess TF binding (Boyle et al. 2008; Buenrostro et al. 2013). DNase-seq and ATAC-seq use proteins – DNase-I and Tn5, respectively – to fragment DNA in open chromatin. TFs bound to accessible DNA disrupt this cleavage, slightly limiting accessibility and leaving a “footprint” of TF binding. These TF footprints can be computationally characterized as small dips in accessibility within regions of high chromatin accessibility (Hesselberth et al. 2009). This technique – digital genomic footprinting – can be used to assay multiple TFs at a time and can provide evidence of in vivo binding events. The increasing availability of ATAC-seq datasets has prompted the development of footprinting tools like HINT-ATAC (Li et al. 2019) and TOBIAS (Bentsen et al. 2020). Though they carry their own caveats, like cleavage biases from Dnase-I and Tn5 and false negatives from transient or weak TF-DNA interactions (Sung et al. 2016; Li et al. 2019; Yan et al. 2020), these tools have proved useful for surveying TF binding in vivo.

By using a combination of measured or predicted TF binding sites, histone marks, chromatin accessibility, high-throughput reporter assay data, and other genome features, computational tools can identify CREs (Lim et al. 2018; Gasperini et al. 2020; Smith et al. 2023). The most accurate models typically rely on many types and/or a large amount of experimental data for training as inputs (Shlyueva et al. 2014; Lim et al. 2018; Smith et al. 2023), which may limit their use in less-characterized cell types or organisms. Given the increasing prevalence of ATAC-seq data, here we wanted to test whether a simple pipeline that primarily relies on ATAC-seq data and digital genomic footprint tools (DGT) could identify novel CREs.

As a test case, we chose to look for CREs active in the *Drosophila melanogaster* segmentation network, which patterns the embryo’s anterior-posterior axis during development. This network is driven by sequentially activated genes encoding TFs, starting with maternal genes, followed by gap genes, pair rule genes, and finally segment polarity genes. This provides a particularly interesting test case to look for novel CREs, as this stage of development has been extensively studied through targeted and genome-wide approaches, yielding a catalog of more than 1,000 characterized CREs (Kvon et al. 2014 June 1; Keränen et al. 2022). To test our approach, we used previously-published ATAC-seq data (Bozek et al. 2019) and the DGF tool TOBIAS (Bentsen et al. 2020) to identify TF footprints in the early *D. melanogaster* embryo genome for 16 essential TFs. We used this data to provide a general survey of where TFs bind in the early embryonic genome. We also used these TF footprints to search for TF clusters that may be novel CREs. With the aid of publicly available histone marks and conservation tracks, we developed a CRE prediction pipeline based on ATAC-seq based DGF. Out of the multiple candidate CREs detected, we isolated five for further functional testing through in situ hybridization and found that all five drove spatially patterned expression, suggesting that this simple pipeline can be applied to other existing ATAC-seq data sets to find new candidate CREs.

## MATERIALS AND METHODS

### ATAC-seq Dataset

We used the ATAC-seq from Bozek et al. (2019), specifically the data from domains D1 and D4, which encompass the anterior and posterior half of the embryo respectively. A whole embryo D1-genotype dataset was used as a control. ATAC-seq reads were mapped in (Bozek et al. 2019) to the dm3 (release 5) genome. Coverage ranged from 50-90 million reads, making it suitable for TF footprint analysis (Yan et al. 2020).

### TF Motif Collection and Testing

TF motifs were retrieved from JASPAR and FlyFactorSurvey (Zhu et al. 2011; Rauluseviciute et al. 2023). Sixteen TFs important in the anterior-posterior patterning of the Drosophila embryo were selected: Bicoid (Bcd), Caudal (Cad), Hb transcription factor (Hb), Giant (Gt), Knirps (Kni), Kr transcription factor (Kr), Tailless (Tll), Huckebein (Hkb), Forkhead (Fkh), Stat92E, Zelda (Zld), Even skipped (Eve), Odd skipped (Odd), Fushi tarazu (Ftz), Hairy (Hry), Runt (Run). Using all reported motifs for each of these 16 TFs (Table S1), we ran TOBIAS v. 0.13.3 BINDetect TF footprint detection with a motif p-value of 0.001 and the dm3 assembly reference genome (Bentsen et al. 2020).

Since many of these TFs had more than one motif reported, to assess which TF motif yielded the most accurate predictions, we compared TF footprints to orthogonal ChIP-seq datasets using the -*intersect* module from the BEDTools suite (Quinlan and Hall 2010). ChIP-seq dataset access was via ReMap2022 (Hammal et al. 2021), and early embryonic stage (stage 4-6) datasets were used (Table S2). We found TF motifs that had a high overlap with ChIP-seq peaks also had the fewest unique hits when contrasted with other motifs for the same TF (Table S3, Figure S1). Therefore, for TFs that lacked ChIP-seq data, the best performing motif was defined as the one that provided the fewest unique predicted binding sites (Table S1). TF footprints for each TF were obtained for D1 and D4 replicates and compiled into a list which was sorted for uniqueness based on genomic coordinates, keeping only the first instance of a TF footprint and removing duplicates.

### TF footprint per Genomic Feature Categorization

TF footprint categorization into different genomic features was done using the -*intersect* module from the BEDTools suite (Quinlan and Hall 2010). The genomic features of interest included known CREs (enhancers), promoters, coding regions (CDS), UTRs, introns, and intergenic regions. The list of known enhancers was manually curated from the REDfly and Fly Enhancers @ Stark Lab Vienna Tile (VT) databases (Kvon et al. 2014 June 1; Keränen et al. 2022).

Enhancers collected from the REDfly database were those with positive expression and active in the blastoderm embryo stage and stages 4-6. Entries belonging to the dm6 assembly of the genome were translated to their dm3 coordinates using the UCSC LiftOver tool (https://genome.ucsc.edu/cgi-bin/hgLiftOver). VTs that were active in the embryonic stages 4-6 but were not present in the REDfly search were added. A union of these two databases provided a collection of 1,232 CRMs acting in the *Drosophila* early embryonic stage.

The partition of the rest of the genome was based on the UCSC dmel r5.57 GFF files (https://ftp.flybase.net/genomes/Drosophila_melanogaster/dmel_r5.57_FB2014_03/gff/). Intergenic regions were defined as regions beyond UTRs, exons, and introns. TF footprints in intergenic regions were collected with the -v option from *bedtools intersect*. Total base pair count per genomic feature was calculated by first merging regions that overlapped into a single region to avoid double counting. Intergenic regions were measured by subtracting the combined base pair count of UTR, exon, and introns from the working genome (without mitochondrial genome and chromosomes U and Uextra)

Promoters were defined as -250 and +50 of from the midpoint of annotated transcription start sites (TSS) (Ohler et al. 2002; Qi et al. 2022). Embryonic RAMPAGE data (Batut and Gingeras 2017) and annotated TSS in the dmel r5.57 genome were considered as candidate TSS definitions. Since RAMPAGE-derived promoters contain a significantly higher number of TF footprints than annotated TSS-derived promoters while maintaining high TF footprint density (Figure S2), we used RAMPAGE-TSS to derive our promoters in the genome-wide TF footprint pattern analysis.

### Candidate Cis-regulatory Element (cCRE) Search

To derive parameters from TF footprint patterns binding within known enhancers, we measured the median TF footprint count and average median TF footprint spacing using custom scripts and the BEDTools *-spacing* module. Initial TF footprint clusters were defined by groups of at least 14 TF footprints in close proximity (11 bp). However, this search yielded no TF footprint clusters beyond known enhancers. Stringency was therefore relaxed to 7 TF footprints at 18 bp apart. TF footprint clusters were overlapped with ChIP-seq peaks of histone marks associated with active enhancers: H3K4me1 and H3K27ac (Creyghton et al. 2010; Koenecke et al. 2017). Enhancer histone mark ChIP-seq data was retrieved from (Li et al. 2014), based on nc14 embryos. Conservation data was retrieved in the form of a track from UCSC PhastCons 15-way track (conservation across 15 insects species: 12 flies, mosquito, beetle, and honeybee) (https://genome.ucsc.edu/cgi-bin/hgc?hgsid=1754389848_7hHSN6x0dBeqQMKAP6xAEIkmYaZ D&db=dm3&c=chr3R&l=26669270&r=26671386&o=26669270&t=26671386&g=phastCons15w ay&i=phastCons15way). Clusters that had more than a 70% overlap with the conservation track were defined as candidate CREs (cCREs).

After gene assignment and identification of cCREs whose target genes expressed distinct spatial patterns, the initial TF footprint cluster region was expanded to cover the entire ATAC-seq peak. These expanded cCREs, now termed high confidence cCREs (hCREs) were chosen for functional assay along with a 600 bp accessible region with no predicted TF footprints.

### cCRE Target Gene Assignment

cCRE target genes were manually selected using a genome viewer. All genes whose TSS were available in a 15kb window flanking highly conserved cCREs were considered possible targets (Arnold et al. 2013). Each candidate target gene was screened for expression in the blastoderm embryo stages (2-4 hrs) based on developmental stage expression summary from FlyBase (Öztürk-Çolak et al. 2024). Each gene expressed at these stages was then filtered and prioritized for body segmentation expression throughout the embryo using the BDGP *in situ* (https://insitu.fruitfly.org/cgi-bin/ex/insitu.pl) (Tomancak et al. 2002) and Fly-FISH databases (https://fly-fish.ccbr.utoronto.ca/) (Lécuyer et al. 2007).

### Cloning and Functional Testing of High Confidence Candidate CREs

Five hCREs and an additional footprint-negative region of open chromatin were cloned from the iso1 Drosophila line. The additional open region is a 600 bp genomic stretch within an ATAC-seq peak that does not possess any predicted TF footprints (Figure S3). The primers used for extraction can be found in Table S4. The six regions were cloned upstream of a minimal Drosophila Synthetic Core Promoter (DSCP) (Pfeiffer et al. 2008) driving the expression of MS2 aptamer sequences and the yellow gene. A mini-white marker was included for selection. Transgenic flies with phi-C31 mediated integration of all six constructs into y[1] w[1118]; PBac{y[+]-attP-3B}VK00002 were generated through BestGene Services. Flies were then crossed to generate hCRE homozygous reporter lines.

### Embryo fixation and HCR v3.0 in situ hybridization

Embryos were collected from population cages after 1 hour of lay time, aged to the developmental stage 5 (nc14), and fixed according to (Keränen et al. 2006). Briefly, after dechorionation in 50% bleach for 3 min, embryos were fixed for 25 min in heptane/10% methanol-free formaldehyde, devitellinized in methanol, washed in methanol and ethanol, and stored in ethanol at −20 °C. Whole-mount RNA fluorescence in situ hybridization was then performed using HCR v3.0 according to the Molecular Instruments protocol for fruit fly embryos. Split-initator probe sets were designed using HCRProbeDesign (Goff lab GitHub) and purchased from IDT. HCRv3 hairpins were purchased from Molecular Instruments. In accordance with the protocol, fixed embryos were hybridized overnight at 37 °C with split-initiator probe sets. A B1 probe set recognizing the *yellow* reporter was included in all samples, together with B5 probe sets for the predicted target genes *brat*, *Svil*, *RhoGAP71E*, *Ac78C*, or *cic*. HCR amplification was performed overnight at room temperature, with amplifier v3 B1-488 for the *yellow* channel and amplifier v3 B5-647 for the target-gene channel.

### Imaging and image processing

Tape spacers were placed between the slide and coverslip to prevent compression and preserve embryo morphology during image acquisition. Whole-mount embryos were mounted in 50% glycerol, 20 mM Tris pH 8.0, and 0.5% N-propyl gallate. Embryos were imaged on a Nikon AX confocal microscope using 488 nm and 647 nm laser lines to detect the *yellow* reporter and endogenous target gene HCR signals, respectively. For each embryo, a single z-stack comprising 9-14 optical sections spaced 5–7 µm apart was acquired, depending on embryo size and orientation. Two acquisition settings were used. Under the higher-power setting, the 647 nm channel was acquired at laser power 12.7 and gain 10, and the 488 nm channel at laser power 19 and gain 10. Under the lower-power setting, the 647 nm channel was acquired at laser power 1 and gain 15, and the 488 nm channel at laser power 1.5 and gain 17. The lower-power setting was used for *RhoGAP71E* and *Ac78C*. Z-stacks were converted to maximum-intensity projections in ImageJ, false-colored using green for the 488 nm channel and magenta for the 647 nm channel.

## RESULTS

### The majority of TF footprints lie outside of characterized CREs

This study aimed to characterize the TF binding site landscape across the *D. melanogaster* early embryo genome using an ATAC-seq-based DGF tool. To do this, we coupled publicly available ATAC-seq data from the anterior (D1) and posterior (D4) regions of nuclear cycle (nc) 14 embryos (Bozek et al. 2019) with TOBIAS, a digital genomic footprinting tool for the prediction of TF binding sites (Bentsen et al. 2020) (Figure 1A). We predicted the binding sites of 16 TFs that play significant roles in anterior-posterior and pair-rule embryonic patterning (Methods). To select among the multiple TF motifs for these factors, we prioritized motifs that either showed the best overlap with ChIP-seq data, or, for those factors without ChIP-seq data, the motif that yielded the fewest unique predicted binding site footprints (Methods). TOBIAS revealed a total of 53,774 footprints for all 16 TFs throughout the genome.

**Figure 1.**
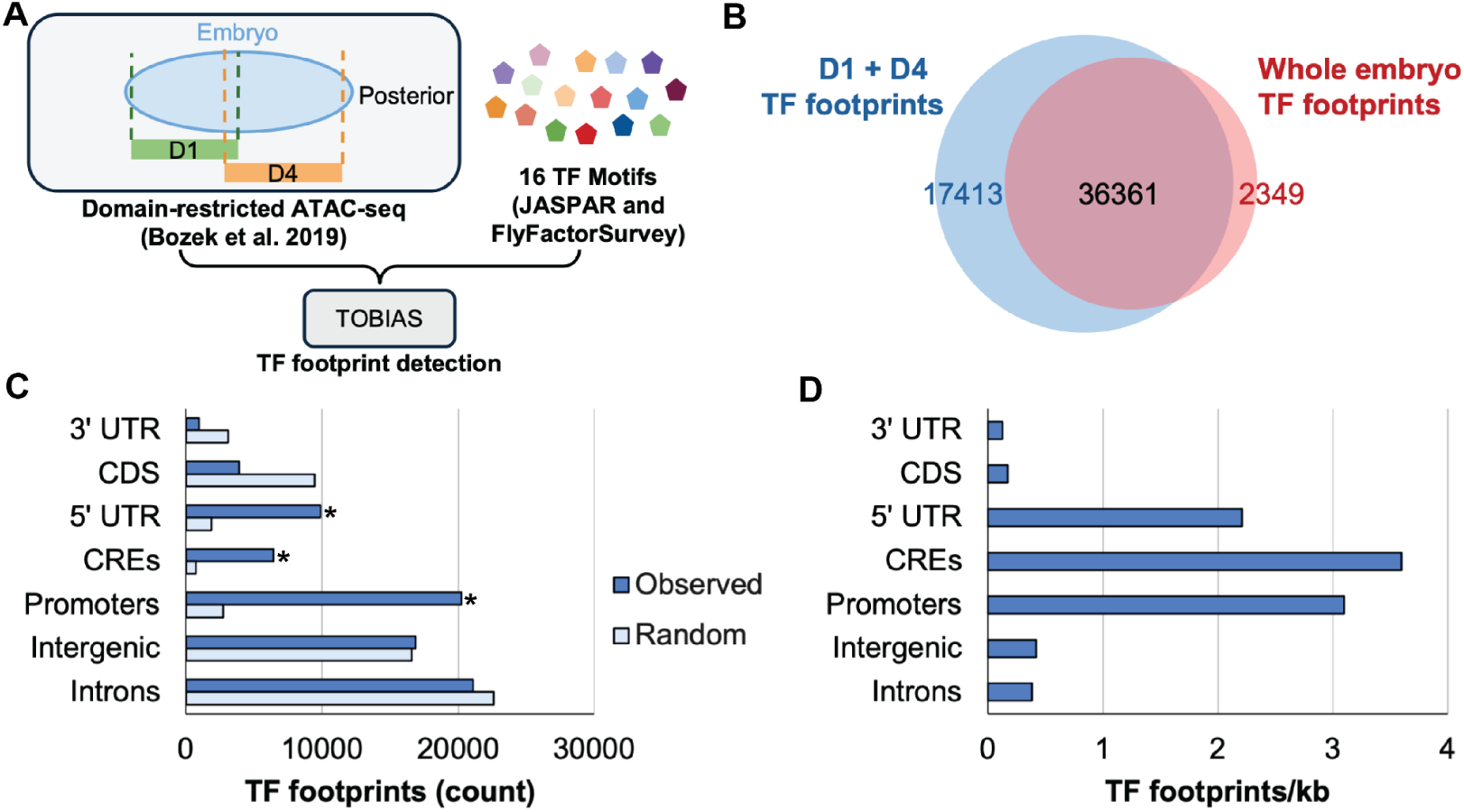
TF footprinting yields a high density of sites in known CREs and a large number of sites outside of annotated regulatory elements. A. ATAC-seq data from the anterior (D1) and posterior (D4) regions of the nc14 *Drosophila* embryo from Bozek et al. (2019) study and 16 TF motifs were used with the TOBIAS digital footprinting tool to detect TF binding events. B. A Venn diagram comparing overlap between TF footprints detected in domain restricted ATAC-seq (D1 + D4) and whole embryo ATAC-seq shows that the regional ATAC-seq data yields a larger number of footprints than the whole embryo data. C-D. The distribution of TF footprints per genomic feature as absolute counts (C) and density (D) is shown. Light blue bars in C indicate the random expectation for TF footprints in each genomic feature, based on the total length in nucleotides of each genomic feature bin. The distribution of TF footprints in each feature differs significantly from the random expectation (*X*^2^ test, *p* < 1e-4), with * indicating genomic features enriched in TF footprints (*X*^2^ post-hoc test with Bonferroni correction, p < 0.05).

Since the TFs expressed and enhancers active in the embryo vary spatially, Bozek, et al. performed ATAC-seq using nuclei from specific regions of the embryo and included a whole embryo ATAC-seq data set as a comparison. Since most of our 16 TFs are not ubiquitously expressed, we hypothesized that the regional ATAC-seq data sets (D1 and D4) would allow us to find TF footprints that are undetectable in the whole embryo dataset, where the footprints might be lost because the sample includes nuclei in which the TF is not expressed. To test this hypothesis, we compared the TF footprint from the regional ATAC-seq with those derived from a whole embryo dataset and predicted the regional data set would yield a larger number TF footprints and a high overlap with whole embryo derived TF footprints. This is indeed the case; 68% (n = 36,361) of the regional-derived TF footprints overlap with the whole embryo-derived predictions (n = 38,710), while 32% (n = 17,413) are unique hits. Additionally, 94% of the whole embryo-derived TF footprints overlap with the regional-derived TF footprints (Figure 1B). These results confirm that ATAC-seq data derived from a more similar population of nuclei, as compared to whole embryo data, allows the identification of a larger number of TF footprints and that footprints found in whole embryo data largely overlap with the regional-derived footprints.

Given the extensive characterization of anterior-posterior patterning enhancers, both in individual studies and high-throughput assessments, we hypothesized a large number of TF footprints would lie in annotated CREs. We searched for overlaps between these footprints and of 1,232 CREs compiled from the REDfly and the Stark Lab Vienna Tile databases (Kvon et al. 2014 June 1; Keränen et al. 2022) (Methods). We found that only around 11.8% of TF footprints overlapped with these regulatory regions (Figure 1C). To understand where the remaining 88.2% of TF footprints are, we categorized the footprints’ location into 5’ UTRs, 3’ UTRs, promoters, coding sequences (CDS), introns, and intergenic regions. This genome-wide survey showed that most TF footprints are in promoters, introns, and intergenic regions. Promoters, CREs, and 5’ UTRs have the highest density of TF footprints (Figure 1D) and are significantly enriched with TF footprints (*X*^2^ test, *p* < 1e-4, post-hoc test with Bonferroni correction, p < 0.05). This analysis confirms that promoters and characterized CREs have high densities of TF footprints, in line with expectation. It also suggests that 5’ UTRs have a high density of TF footprints, which we confirmed was not simply due to overlapping promoter/5’ UTR annotations (Figures S4 and S5). Lastly, the large absolute number of TF footprints outside of known regulatory elements suggest that there may be many unannotated CREs.

### TF footprint clusters may be uncharacterized CREs

We noted that there was a high frequency, albeit low density, of TF footprints in intergenic and intronic regions. We hypothesized TF footprint clusters in these regions may be uncharacterized CREs. To address this question, we measured the number of TF footprints and the spacing between them in known enhancers. Since some of the known enhancers may not be involved with anterior-posterior patterning or bound by the analyzed TFs, we first analyzed known CREs with at least one TF footprint and found they had a median of 14 TF footprints per region (Figure 2A), spaced at a median of 11 bp apart (Figure 2B). To find potential novel CREs, we searched for TF footprint clusters that also shared these features and found no additional TF footprint clusters. We therefore relaxed the search criteria to 7 TF footprints per cluster, based on our calculated median across all collected CREs and a a previous report of the average TF binding site composition per CRE (Yang et al. 2022), and increased the maximum spacing between them to 18 bp. This search yielded 450 TF footprint clusters. However, we expected that some of these clusters might not have regulatory potential, since TF binding is not always indicative of regulatory activity (Li et al. 2008; Spivakov 2014).

**Figure 2.**
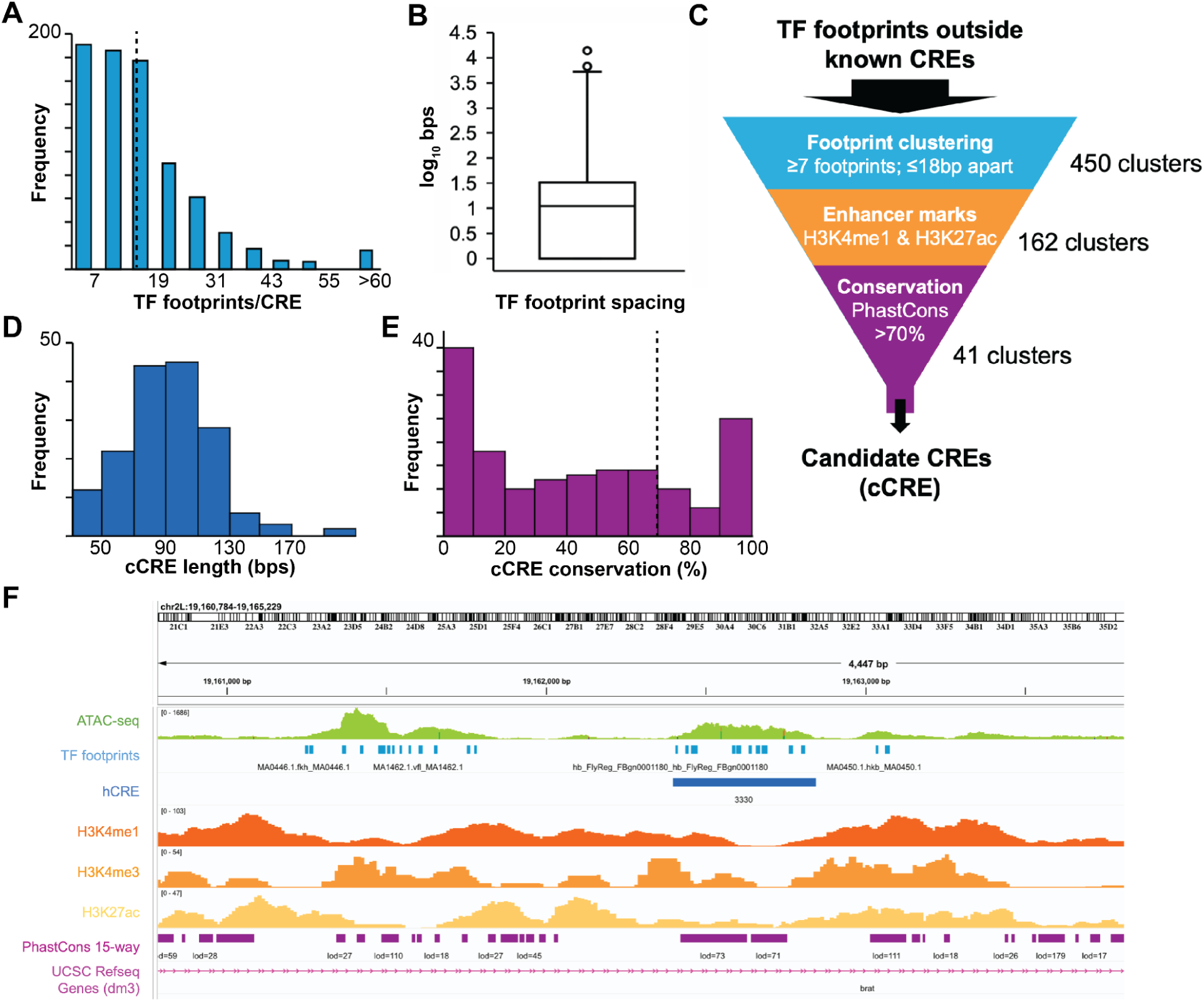
TF footprint patterns in known CREs. **A.** Distribution of TF footprints per known CREs that have at least one TF footprint (n=783). The dashed line indicates the median TF footprint per CRE (14 TF footprints). **B.** Boxplot of TF footprint spacing (log_10_ bps) in known CREs. Median spacing between TF footprint is 11 bp. **C.** TF footprints outside of known CREs were passed through a series of filters. Clusters of TF footprints were tested for their overlap in both active enhancer histone marks (H3K4me1 and H3K27ac) and then filtered by conservation level. **D.** Length distribution of TF footprint clusters that overlap active enhancer marks. Mean number of bp per TF footprint cluster is 93bp. **D.** Histogram of identified TF footprint cluster percent overlap with PhastCons15-way track with the dashed line indicating the 70% overlap threshold. 41 TF footprint clusters are highly conserved. **F.** Example Genome Track View for hCRE 3330 in the Intron of the *brat* gene. Top track indicates ATAC-seq data from Bozek et al. (2019). Active enhancer histone marks H3K4me1 and H3K27ac are displayed together with the promoter histone mark H3K4me3 in track underneath the hCRE track. The conservation information from PhastCons 15-way track and their respective LOD scores are also displayed. Genome track views for the rest of the hCREs can be found in Figure S3.

To narrow down this list to find potential CREs, we screened each cluster for possible regulatory activity using enhancer histone marks and sequence conservation (Figure 2C; (Shlyueva et al. 2014; Lim et al. 2018; Andersson and Sandelin 2020). We identified TF footprint clusters that overlapped with active enhancer H3K4me1 and H3K27ac ChIP-seq peaks (Li et al. 2014). This resulted in a list of 162 clusters with an average size of 93 bp (Figure 2D). We then identified clusters that overlapped with the UCSC PhastCons 15-way track. Out of the 162 clusters, 41 had a conservation of 70% or higher (Figure 2E). We termed these 41 clusters as candidate CREs (cCREs).

To further characterize the regulatory potential of these 41 clusters, we annotated each cluster’s possible gene targets. To assign target genes, we considered any TSS that fell in 15 kb windows flanking each cCREs. Since we were testing TFs important in embryonic patterning, we prioritized cCREs whose potential gene target had a spatially-varying expression pattern throughout the embryo using in situ hybridization data from the BDGP and Fly-FISH databases (Tomancak et al. 2002; Lécuyer et al. 2007). We categorized each cCRE target gene as Patterns Present, Ubiquitous Strong, Ubiquitous, or No Expression. Table S5 shows the results of this annotation. From these 41 clusters, we isolated 5 high confidence CREs (hCREs) (Table 1, Figure 2F). Only one of the hCREs, 16864, overlaps with known CREs: two *cic* CREs (cic_GMR90B07 and cic_HT27), neither of which has been assayed during the early embryonic stage.

**Table 1.**
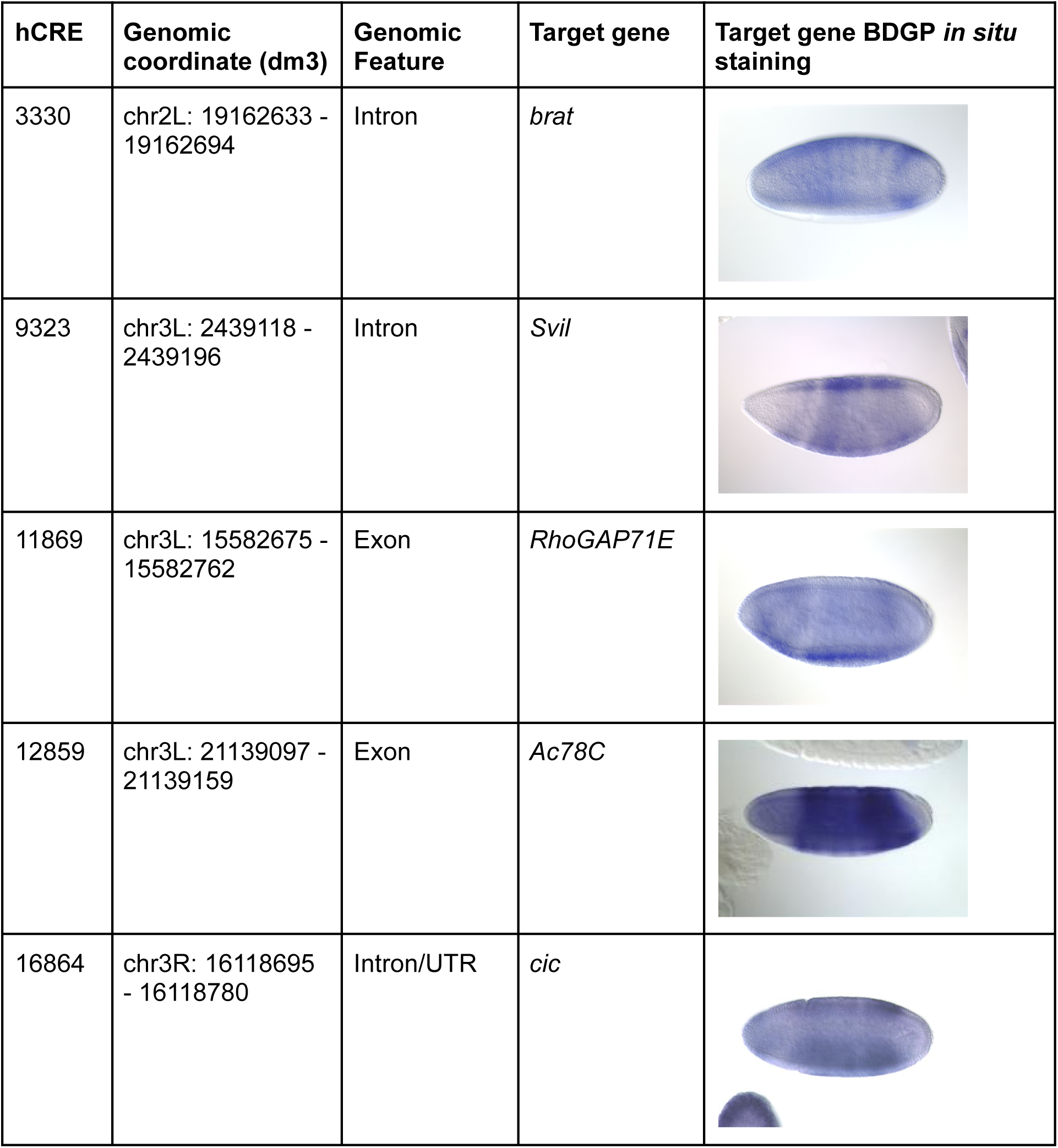
Genomic Features of High Confidence Candidate CREs (hCRE) Putative target genes were assigned based on proximity (within 15kb) to the hCRE and spatial expression patterns informed by the BDGP in situ database.

| hCRE | Genomic coordinate (dm3) | Genomic Feature | Target gene | Target gene BDGP *in situ* staining |
| --- | --- | --- | --- | --- |
| 3330 | chr2L: 19162633 - 19162694 | Intron | *brat* |  |
| 9323 | chr3L: 2439118 - 2439196 | Intron | *Svil* |  |
| 11869 | chr3L: 15582675 - 15582762 | Exon | *RhoGAP71E* |  |
| 12859 | chr3L: 21139097 - 21139159 | Exon | *Ac78C* |  |
| 16864 | chr3R: 16118695 - 16118780 | Intron/UTR | *cic* |  |

### Reporter assays confirm the in vivo activity of hCREs

Previous studies have underscored the importance of functionally testing potential CREs to ascertain their regulatory potential (Halfon et al. 2011). To test our hCREs, we first increased the length of clusters to cover the entire ATAC-seq peak. This expanded our hCREs to genomic fragments ranging from 357 to 978 bp. We compared our expanded hCREs to the PhastCons 15-way track again and found that conservation was still high, with an average of 74% (Table 2). We then cloned these expanded hCREs upstream of a minimal DSCP promoter (Pfeiffer et al. 2008), driving the expression of the *yellow* reporter gene and integrated each into the fly genome at a specific location in the genome. As a point of comparison, we also generated an additional reporter with a 600 bp region that lies in open chromatin, but with no TF footprints.

**Table 2.** Final hCRE Conservation Scores.

| hCRE | TF footprint cluster size (bp) | ATAC-seq peak region (bp) | Peak region % Conservation |
| --- | --- | --- | --- |
| 3330 | 61 | 446 | 71.5 |
| 9323 | 78 | 978 | 67.6 |
| 11869 | 87 | 357 | 82.9 |
| 12859 | 62 | 617 | 77.3 |
| 16864 | 85 | 635 | 68.6 |

hCRE original sizes (TF footprint cluster size) and their final extended sizes (ATAC-seq peak region). Peak region % conservation is based on the overlap between the extended sizes and the PhastCons 15-way region.

To characterize each hCRE’s activity, we used HCR to stain for the yellow reporter gene and the putative target gene of each hCRE. All five hCREs and the region of open chromatin drove yellow expression in a spatially patterned manner, both at stage 5 of development and in various later stages (Figure 3). In most cases, the hCRE activity matched the expression pattern of the putative target gene, except for *brat*, where the hCRE construct drove *brat*-like signal whereas the target stain had no detectable signal in stage 5, and a less distinct pattern of expression in a later stage of development. The footprint-negative region of open chromatin drove a robust, pair-rule-like expression in both stage 5 and later (Figure 3F), suggesting that our hCRE pipeline may yield false negatives.

**Figure 3.**
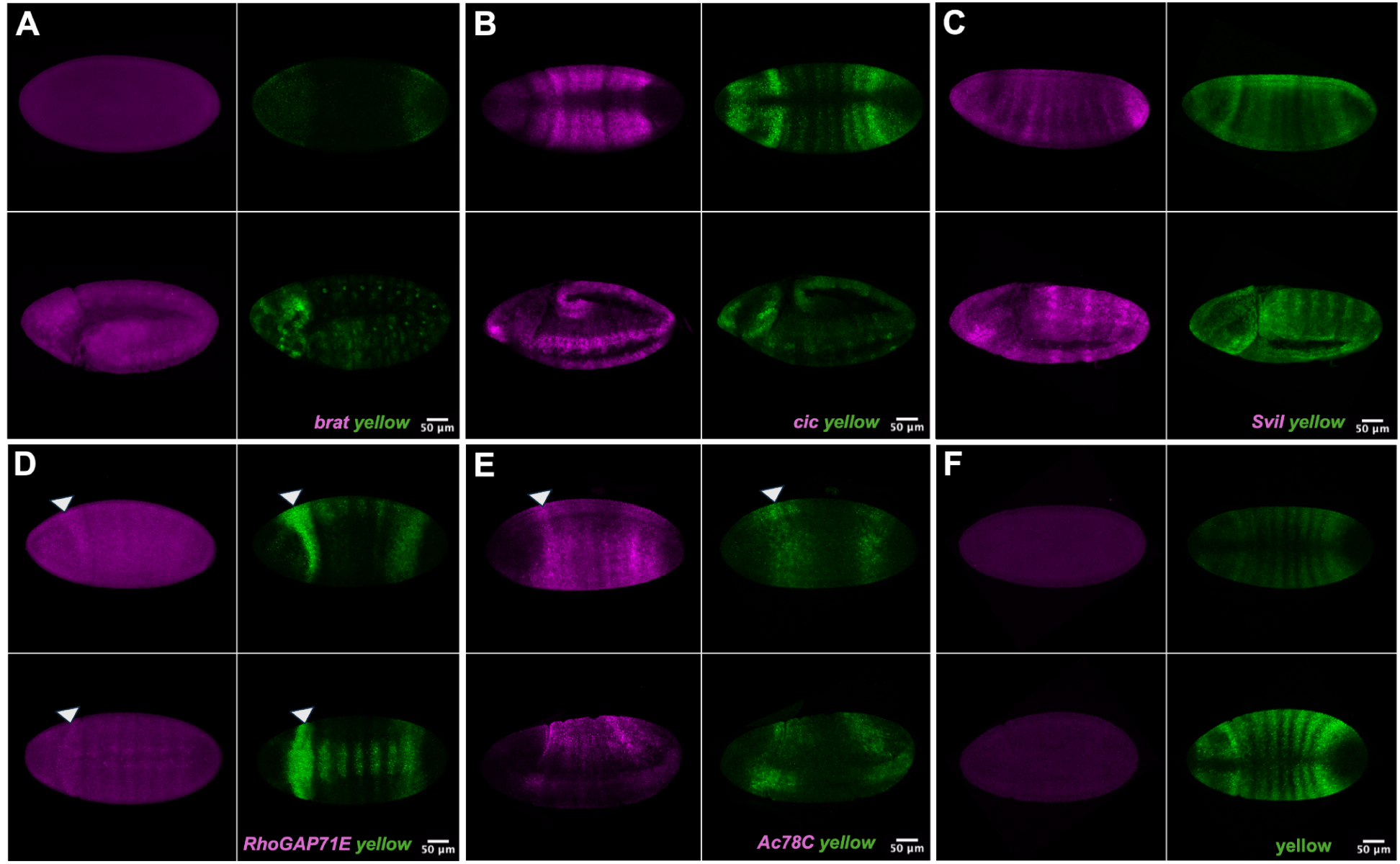
Whole-mount HCR v3.0 detection of endogenous target genes and the *yellow* reporter in *Drosophila* embryos. Representative maximum-intensity projections of embryos processed by whole-mount HCR v3.0. Endogenous target-gene transcripts are shown in magenta (B5-647) and the *yellow* reporter transcript is shown in green (B1-488). For each genotype, the upper image shows a stage 5 embryo and the lower image shows an embryo at post-stage 5. Panels show the following target genes: (A) *brat*, (B) *cic*, (C) *Svil*, (D) *RhoGAP71E*, (E) *Ac78C*, and (F) footprint-negative region stained for *yellow* only.

## DISCUSSION

In this study, we used the DGF tool TOBIAS (Bentsen et al. 2020) to leverage previously published ATAC-seq data of the *D.melanogaster* early embryo (Bozek et al. 2019) to create a genome-wide mapping of anterior-posterior patterning TF footprints. We found a high density of TF footprints in known CREs and promoters, and a large number of footprints outside these regions. Therefore, together with publicly available functional genomic datasets, we constructed a pipeline to identify uncharacterized CREs active in the early embryonic stage of *D.melanogaster* from TF footprint clusters. Functional tests of five novel CREs confirmed that this pipeline can find uncharacterized CREs.

### Digital genomic footprint tools are affected by motif quality and can miss binding events

Many of the TFs we studied had more than one available binding motif (Table S1), and different motifs yielded distinct TF footprint data sets with variable overlaps with ChIP-seq datasets (Table S3). We prioritized motifs with the highest percent overlap with ChIP-seq data, or when lacking stage-matched ChIP-seq data, the motifs with the fewest unique predicted footprints (Methods). For some TFs, namely Zld, the overlap between all motifs and ChIP-seq was quite high, exceeding 70%, while others were much lower (e.g. Bcd, with 36-39% overlaps; Table S3). This may reflect differences in the biochemistry of TF binding – Zld is a pioneer factor and Bcd can bind cooperatively or indirectly, and DGF tools will not detect indirect interactions (Burz et al. 1998; Lebrecht et al. 2005; Schulz et al. 2015; Sun et al. 2015). Additionally, the authors of TOBIAS estimated that ∼40% of TFs do not leave strong footprints (Bentsen et al. 2020).

Together, this suggests that motif quality will impact the accuracy of DGF tools, and that individual binding events may be missed. However, even with these caveats, our results support that searching for clusters of footprints to identify new CREs can still be fruitful. To alleviate the impact of binding events missed by DGF tools, we recommend using a collection of TFs active in the same gene expression patterning process.

### A potential regulatory role of TF footprints in 5’ UTRs

Beyond predicting CREs, we were also interested in better understanding the TF footprint landscape in the early embryo genome. One unexpected observation was the high frequency and density of TF footprints in 5’ UTRs. Our results show that 34% of these 5’ UTR TF footprints do not overlap with promoters, prompting us to ask the functional roles these TF footprints could play (Figure S4; Figure S5). Single nucleotide polymorphism (SNP) variants in these untranslated regions are expected to have an effect on translational regulation, rather than transcriptional control (Ryczek et al. 2023). However, SNPs affecting binding sites of Smad3 in human and mouse 5’ UTR regions can modulate transcriptional regulation (Córdova et al. 2015; Xie et al. 2020). While surveying ChIP-seq peaks in human exonic regions, Chen et al. proposed that TF footprints in 5’ UTRs could potentially enhance the transcription levels of nearby genes (Chen et al. 2023). Our study proposes *D.melanogaster* as another case study where extensive TF footprints in 5’ UTRs are detected, and our observations encourage future work on understanding the role of these binding sites.

### Study limitations

In our study, we identified 41 TF footprint clusters as cCREs using a combination of ATAC-seq TF footprints, histone ChIP-seq, and sequence conservation data with highly stringent thresholds, particularly for conservation. Sequence conservation has been previously used to identify CREs, e.g. (Berman et al. 2002; Halfon et al. 2011; Chen et al. 2018; Moudgil et al. 2023). However, the assumption that CREs are under purifying selection and conserved at that sequence level needs to be taken into careful consideration. The UCSC PhastCons 15-way track includes insects outside of the *Drosophila* clade such as mosquitoes, red flour beetles, and the honeybee (*Anopheles gambiae, Tribolium castaneum,* and *Apis mellifera*), but the gene regulatory network controlling anterior-posterior patterning differs between these species. For example, *bicoid* is present in the *Drosphila* genus and other higher Dipterans, but absent in mosquitos, red flour beetles, and the honeybee (McGregor 2005). Furthermore, gene expression activity can be maintained with extensive CRE sequence divergence or replacement of one CRE with another, and species can harbor unique CREs (Ludwig et al. 2005; Hare et al. 2008; Arnold et al. 2014; Villar et al. 2015). A less stringent conservation threshold or its elimination would expand our list of TF footprint clusters to others that may have regulatory potential.

We note other study limitations. When studying different tissues, histone enhancer mark and target gene expression data may not be available. In these cases, a pipeline relying solely on ATAC-seq TF footprints and, optionally, sequence conversation could be developed, though it is possible that this pipeline would yield false positives. In addition, our footprint-free region of open chromatin drove a distinct pair-rule like pattern of gene expression (Figure 3F), despite the lack of detectable TF binding (Figure S5). There are several possible interpretations of this result. There may be TF binding in vivo that was missed by the TOBIAS algorithm. However, the genes within 15 kb of this region, *mtgo, her, grp, and Trpɣ,* lack distinct spatial expression patterns at this time in development. It is also possible this region can drive expression in the reporter we generated, but not in its endogenous context. Though the region is in open chromatin in the embryo, it is possible that it does not interact with any neighboring genes due to promoter incompatibility or insulator elements. Nevertheless, we find that TF footprinting is a useful way to analyze existing ATAC-seq data to reveal novel cis-regulary elements, even in a relatively well-characterized developmental stage.

## DATA AND CODE AVAILABILITY

All scripts associated with this project and TF footprint prediction BED files can be found in the following GitHub Repository: https://github.com/WunderlichLab/ATAC_footprinting_analysis_project

## STUDY FUNDING

This work was funded by NIH award R01HD095246 to Z.W. and Boston University’s Marian Kramer Award, Multicellular Design Center Kilachand Fellowship, and BU Nano Fellowship to

J.N. C.M. was supported by Boston University’s Undergraduate Research Opportunities Program and the Boston University STEM Pathways Program (Department of Defense DoD STEM FY20 Award HQ00342110008). The content is solely the responsibility of the authors and does not necessarily represent the official views of any of the funders.

## CONFLICTS OF INTEREST

The authors declare no conflicts of interest.

## Supplementary Tables and Figures

**Table S1.** Transcription Factor Binding Motifs. Collection of all TF motifs accessed from JASPAR and FlyFactorSurvey. Motifs marked in green were chosen for the downstream analysis.

| Transcription Factor | JASPAR Motifs | FlyFactorSurvey Motifs |
| --- | --- | --- |
| Bicoid | MA0212.1 | Bcd_SOLEXA, Bcd_NAR, Bcd_Cell, bcd (bcd_FlyReg_DnaseI-data) |
| Caudal | MA0216.1, MA0216.2 | Cad_SOLEXA, Cad_Cell, cad(cad_FlyReg_DnaseI-data) |
| Giant | MA0447.1 | gt_NAR, gt (gt_FlyReg_DnaseI-data) |
| Hb transcription factor | MA0049.1 | hb_NAR, hb_SANGER, hb_SOLEXA, hb (hb_FlyReg_DnaseI-data) |
| Knirps | MA0451.1, MA0451.2 | kni_SANGER, kni (kni_FlyReg_DnaseI-data) |
| Kr transcription factor | MA0452.1, MA0452.2, MA0452.3 | Kr_SANGER, Kr_SOLEXA, Kr (Kr_FlyReg_DnaseI-data) |
| Stat92E | MA0532.1, MA0532.2 |  |
| Tailless | MA0459.1 | tll (tll_FlyReg_DnaseI-data) |
| Zelda (vlf) | MA1462.1, MA1462.2 | vlf_SANGER, vlf_SOLEXA |
| Huckebein | MA0450.1 |  |
| Forkhead | MA0446.1 |  |
| Even skipped | MA0221.1 | Eve_SOLEXA, Eve_Cell, eve (eve_FlyReg_DnaseI-data) |
| Odd skipped | MA0454.1 |  |
| Fushi tarazu | MA0225.1 | Ftz_Cell, Ftz_SOLEXA, ftz (ftz_FlyReg_DnaseI-data) |
| Hairy | MA0449.1 | h_SANGER |
| Runt | MA2319.1 | run_Bgb_NBT |

**Table S2.** ChIP-seq Data Sets. Gene Expression Omnibus (GEO) accession numbers for ChIP-seq data sets used to compare TOBIAS binding site predictions to in vivo binding and overlap TF footprints with enhancer histone marks.

| **Transcription Factor/Histon Mark** | **ChIP-seq GEO accession number** |
| --- | --- |
| Bicoid | GSE86966 |
| Zelda | GSM763062 |
| Giant | GSE50771 |
| Hb transcription factor | GSE50771 |
| H3K4me1 | GSE58935 |
| H3K27ac | GSE58935 |

**Table S3.** Amount of overlap between TOBIAS-predicted binding sites and ChIP-seq data for four TFs. Overlap between TOBIAS predicted TF footprint from different TF motifs and corresponding ChIP-seq peak

| **Bcd motif** | **% overlap with ChIP-seq peaks (GSE86966)** |
| --- | --- |
| MA0212.1 | 37 |
| Bcd_SOLEXA | 37 |
| Bcd_NAR | 37 |
| Bcd_Cell | 36.5 |
| bcd (bcd_FlyReg_DnaseI-data) | 39 |
| **Zld motif** | **% overlap with ChIP-seq peaks (GSM763062)** |
| MA1462.1 | 78 |
| MA1462.2 | 79 |
| vlf_SANGER | 75 |
| vlf_SOLEXA | 73 |
| **Gt motif** | **% overlap with ChIP-seq peaks (GSE50771)** |
| MA0447.1 | 12.9 |
| gt_NAR, | 12.4 |
| gt (gt_FlyReg_DnaseI-data) | 9.5 |
| **Hb motif** | **% overlap with ChIP-seq peaks (GSE50771)** |
| MA0049.1 | 8.2 |
| hb_NAR | 6.6 |
| hb_SANGER | 7.3 |
| hb_SOLEXA | 7.8 |
| hb (hb_FlyReg_DnaseI-data) | 8.2 |

**Table S4.**
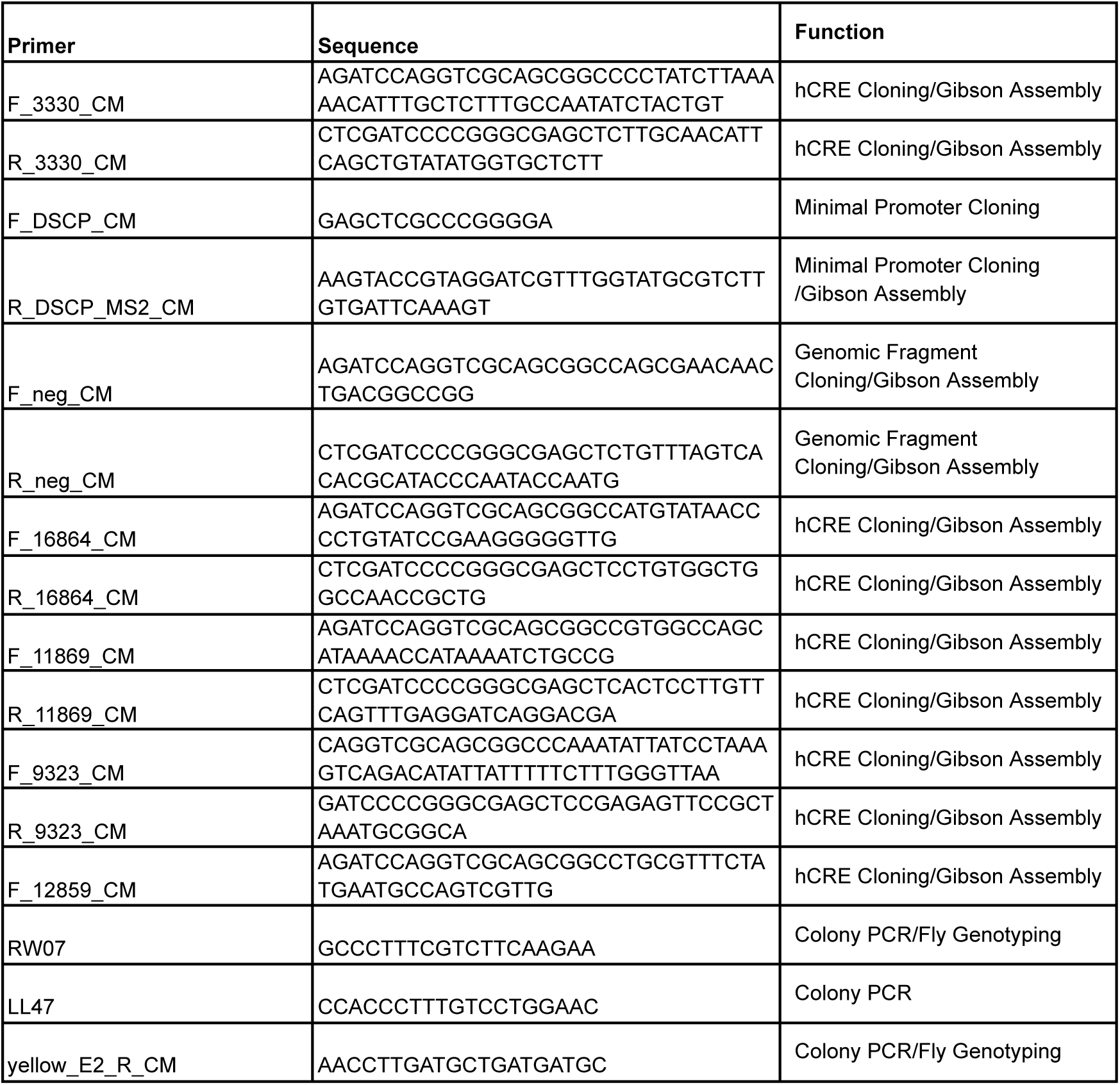
Primers used in this study. Primers for genomic extraction of hCREs, isolation of the DSCP minimal promoter, subsequent colony PCRs and fly genotyping.

**Table S5.**
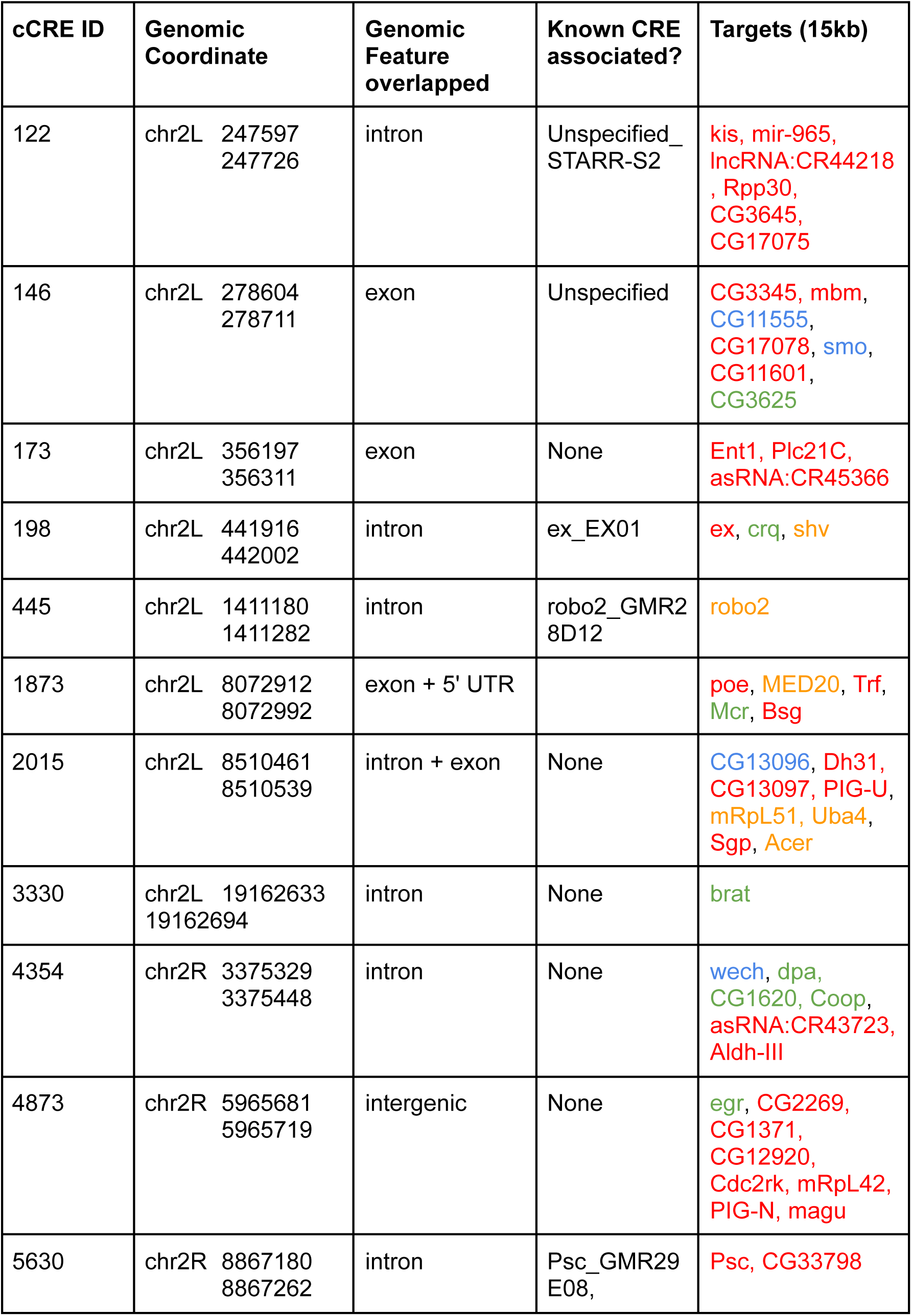
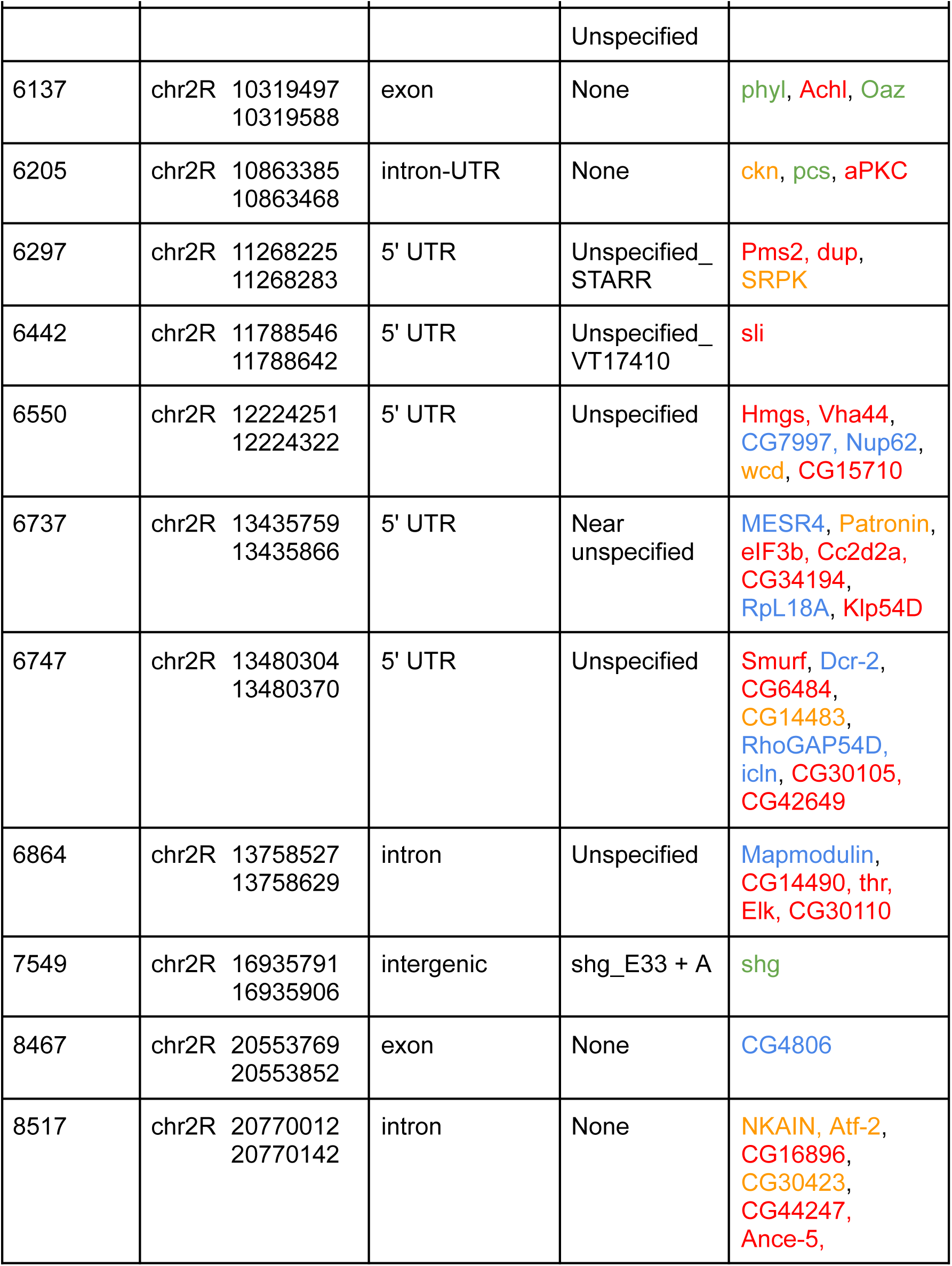
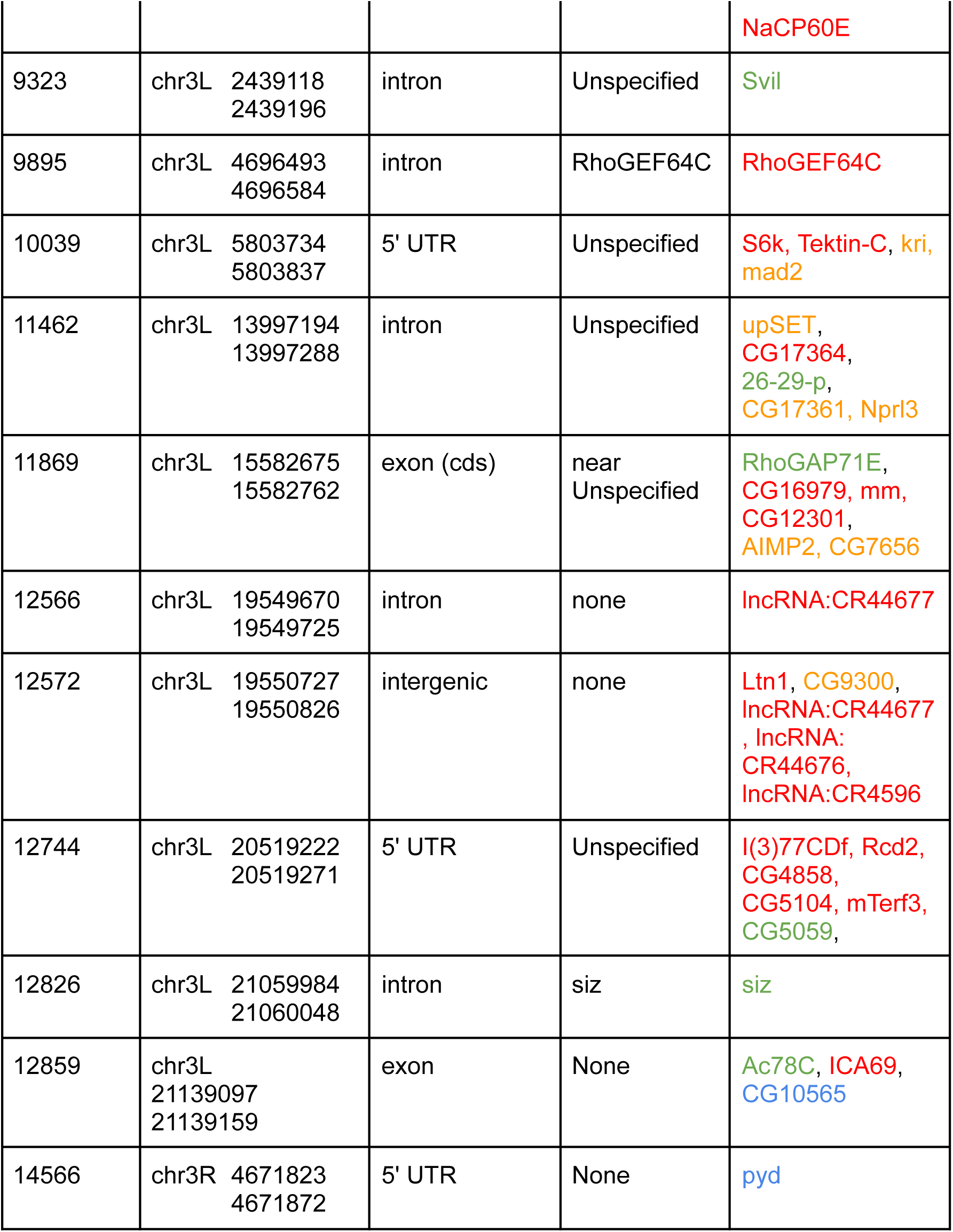
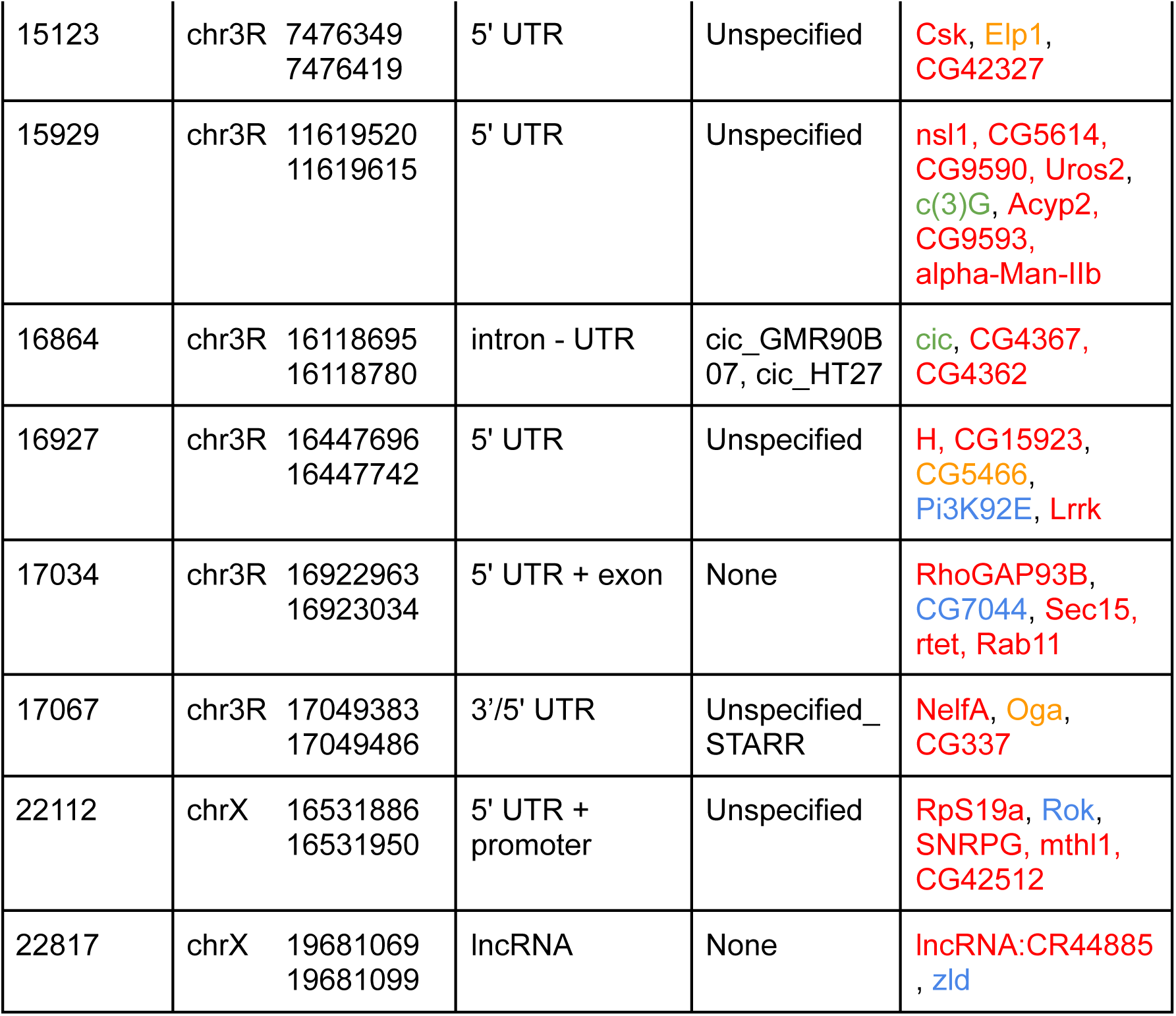
cCRE Clusters and Their Target Genes.

**Figure S1.**
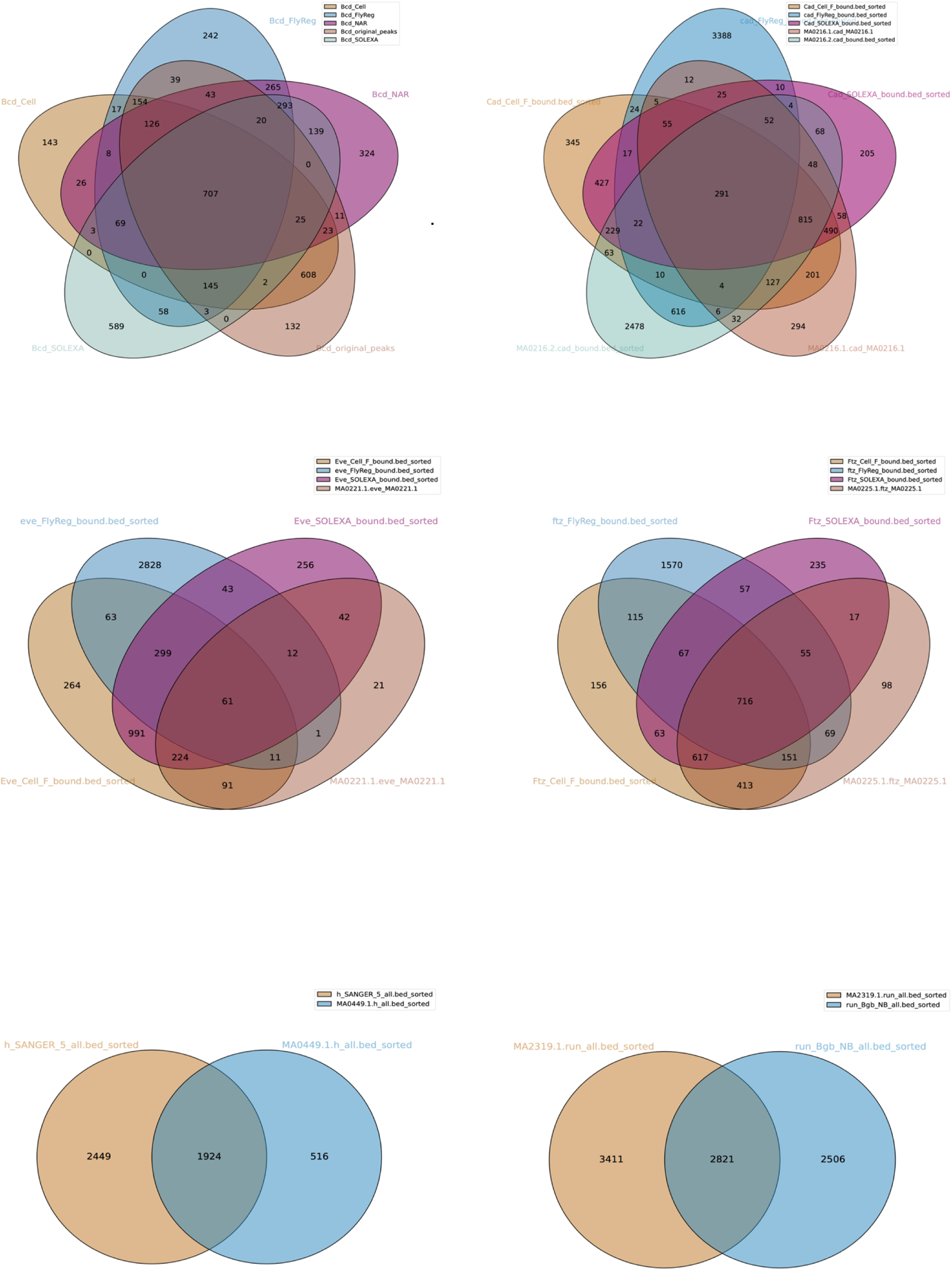
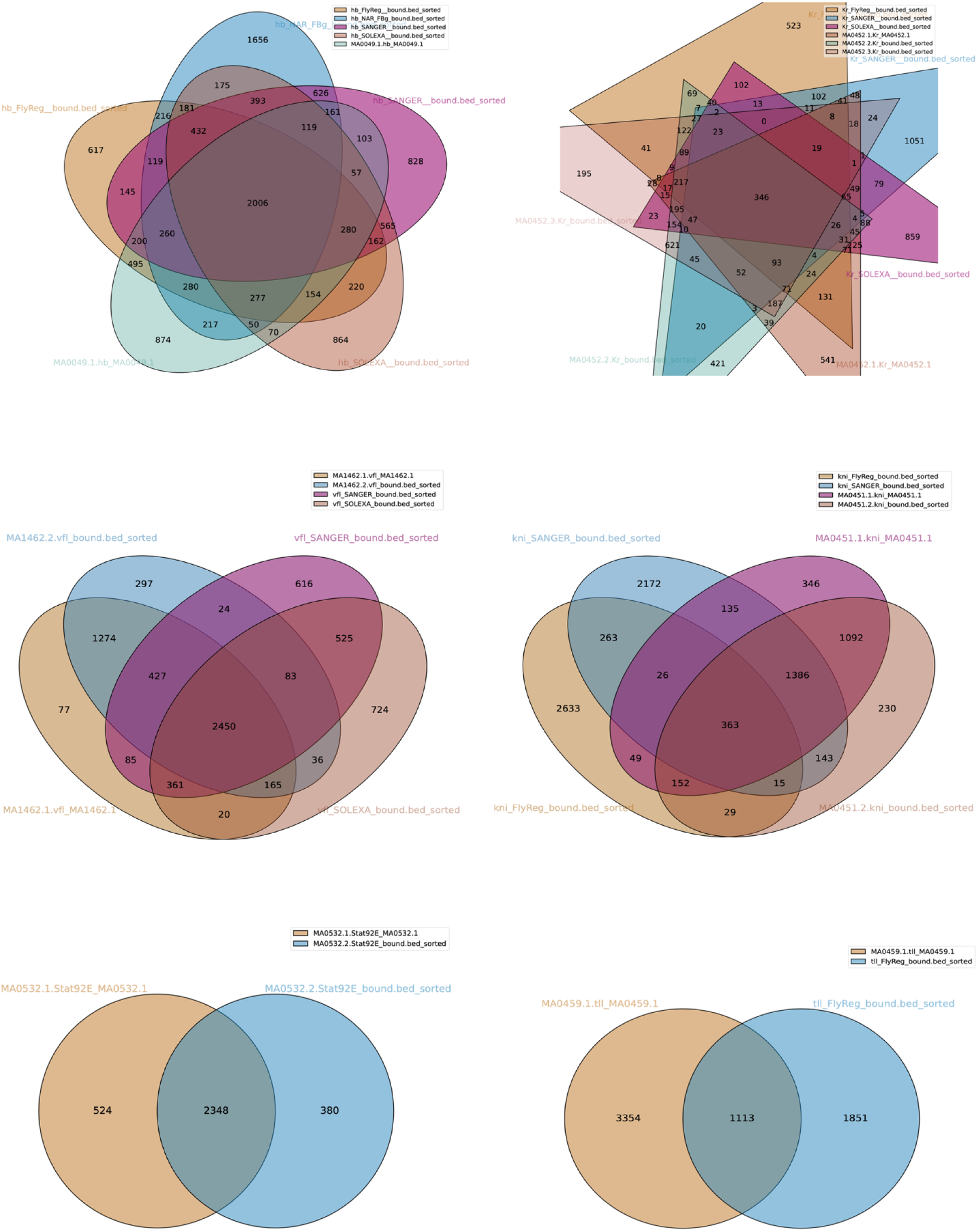
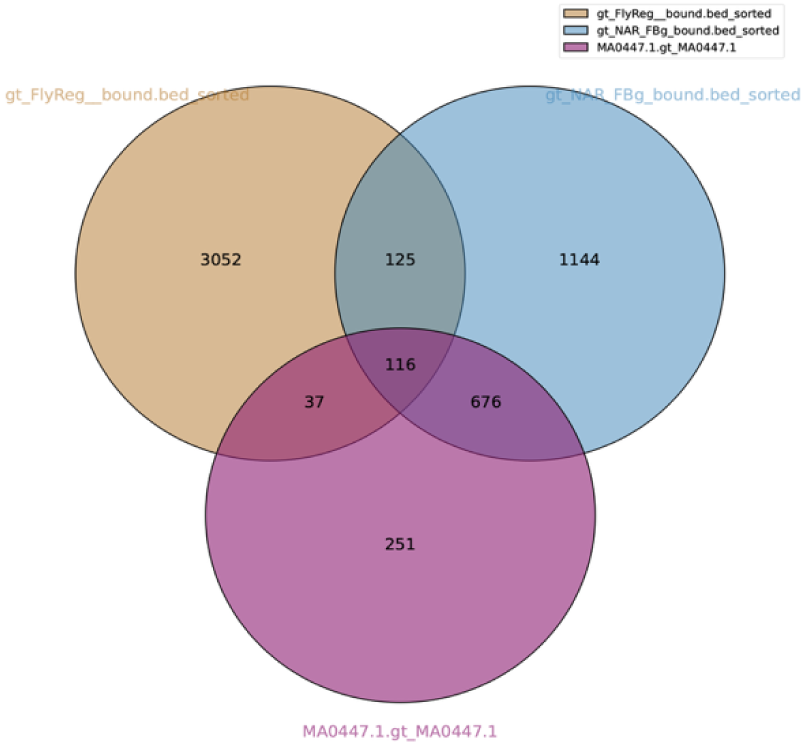
Overlaps between TOBIAS-predicted sites for different TF binding motifs. Venn diagrams comparing predicted outputs from different TF motifs. Bcd_original_peaks are TF footprint derived from JASPAR’s MA0212.1 Bcd motif, and vlf is the old gene symbol for Zelda. Motifs with the fewest unique predicted binding sites were retained for downstream analysis (Table S1).

**Figure S2.**
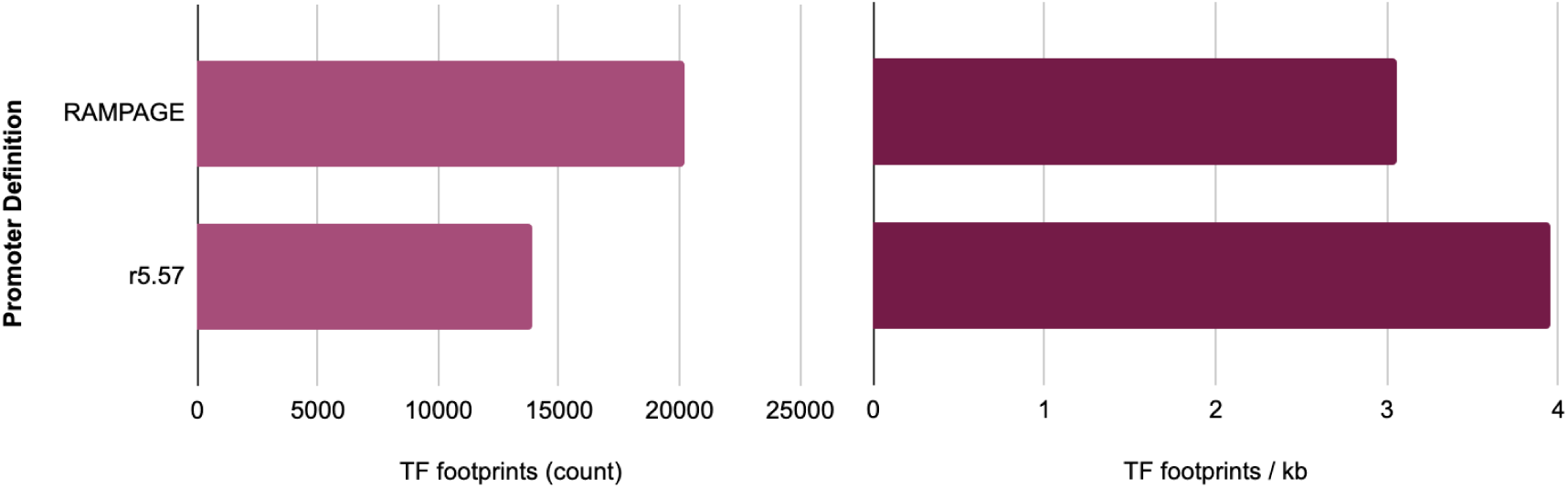
RAMPAGE-based Promoter Definition Captures More TF footprints. TF footprint count (left) and density (right) by promoter definition. Every promoter region was-250 and +50 from the midpoint of the TSS. TSS were mined from dm3 r5.57 GFF file or from RAMPAGE data (Batut and Gingeras 2017).

**Figure S3.**
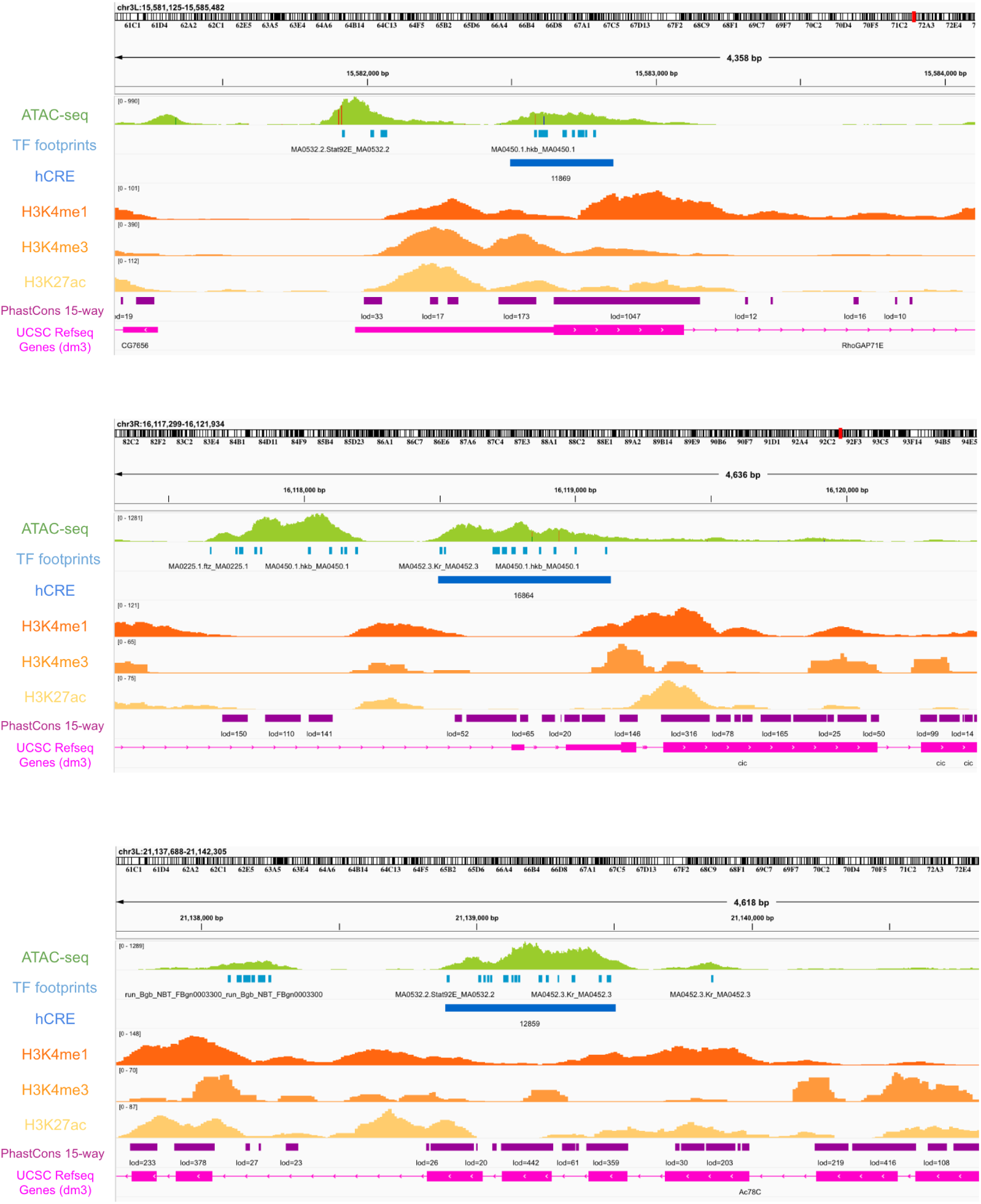
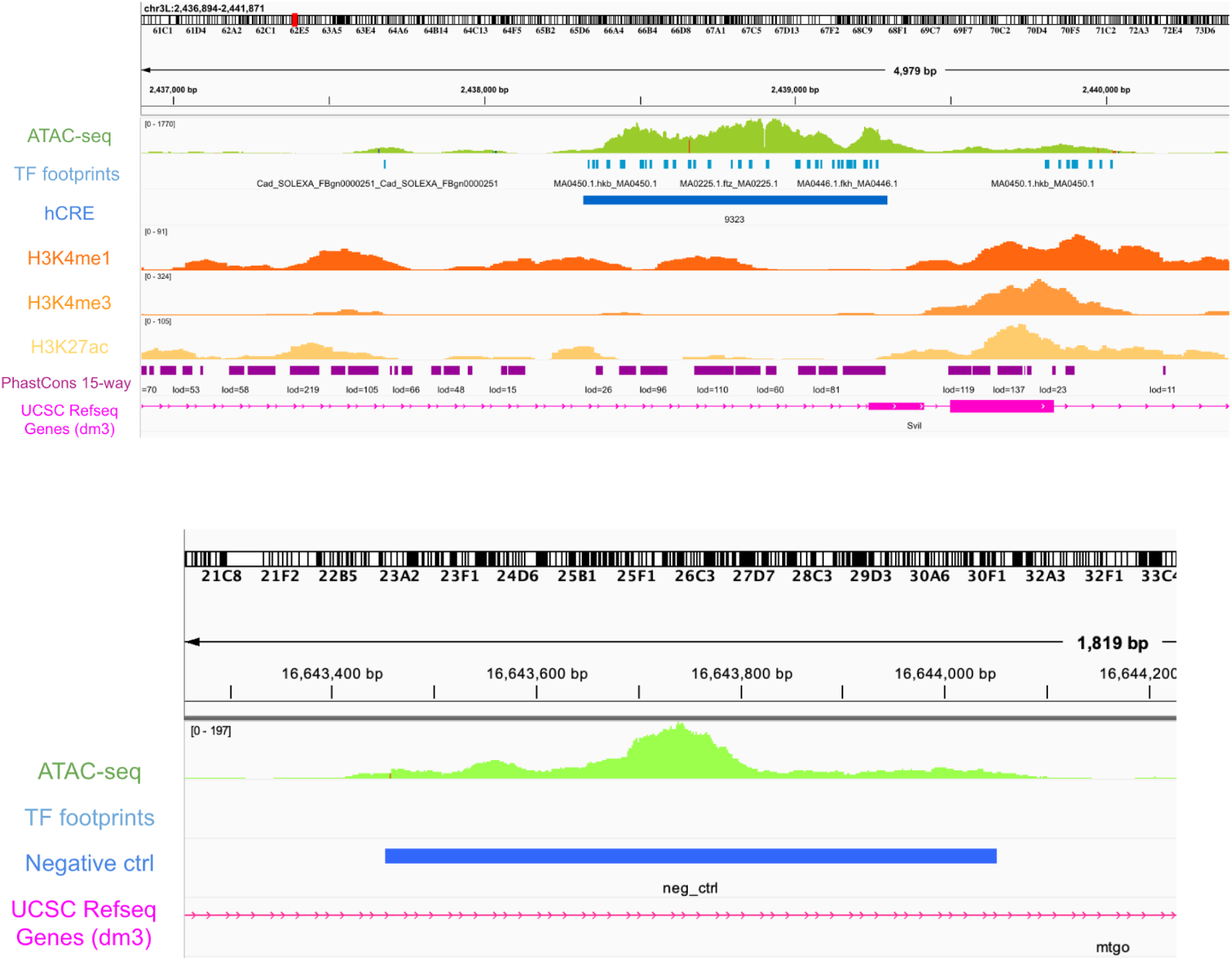
Genome Browser views for hCREs and the footprint-negative region cloned into reporter constructs. For the hCREs, top track indicates ATAC-seq data from Bozek et al. (2019). Active enhancer histone marks H3K4me1 and H3K27ac are displayed together with the promoter histone mark H3K4me3 in track underneath the hCRE track. The conservation information from PhastCons 15-way track and their respective LOD scores are also displayed. The target gene is indicated in the UCSC Refseq Gene track. The footprint-negative region is accessible in the embryonic genome as informed by ATAC-seq data but did not have any TF footprints.

**Figure S4.**
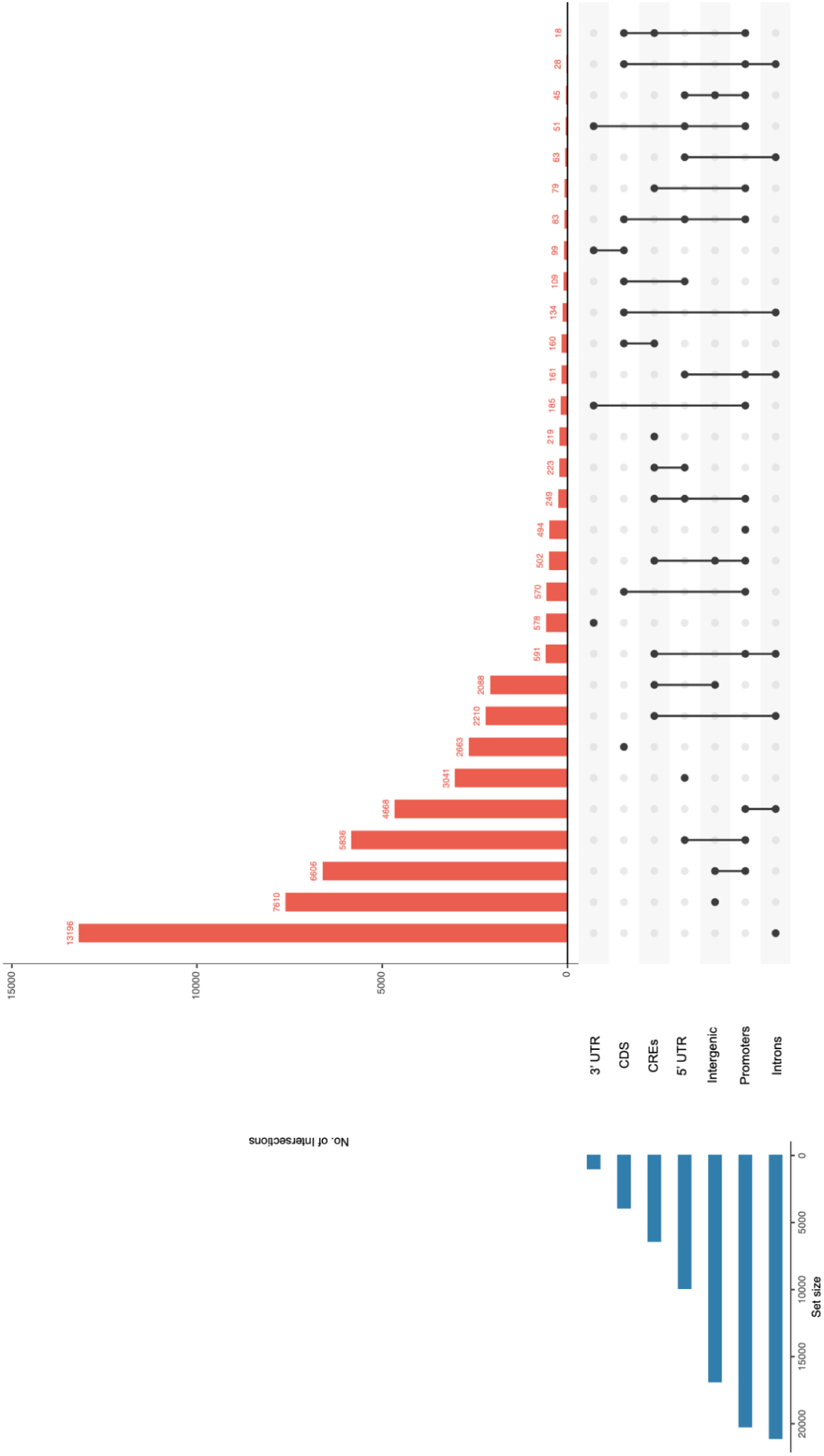
Intersection of TF footprints per genomic feature. Intervene UpSet graph of all TF footprints per genomic features to show TF footprints that bind to two or more genomic features. Bar chart on the lower left shows the frequency of TF footprint per genomic feature. Dots on the right indicate which datasets are part of the intersection. Single dots are TF footprints unique to the set, and connected dots indicate groups of TF footprints that overlap in different genomic features. Frequency of these overlaps are displayed in the bar chart on top. These results show that ∼66% of TF footprints in 5’ UTRs overlap with TF footprints found in promoters, while the remaining 34% are unique to 5’ UTRs, suggesting that there are a number of TF footprints in 5’ UTRs that do not overlap with promoters.

**Figure S5.**
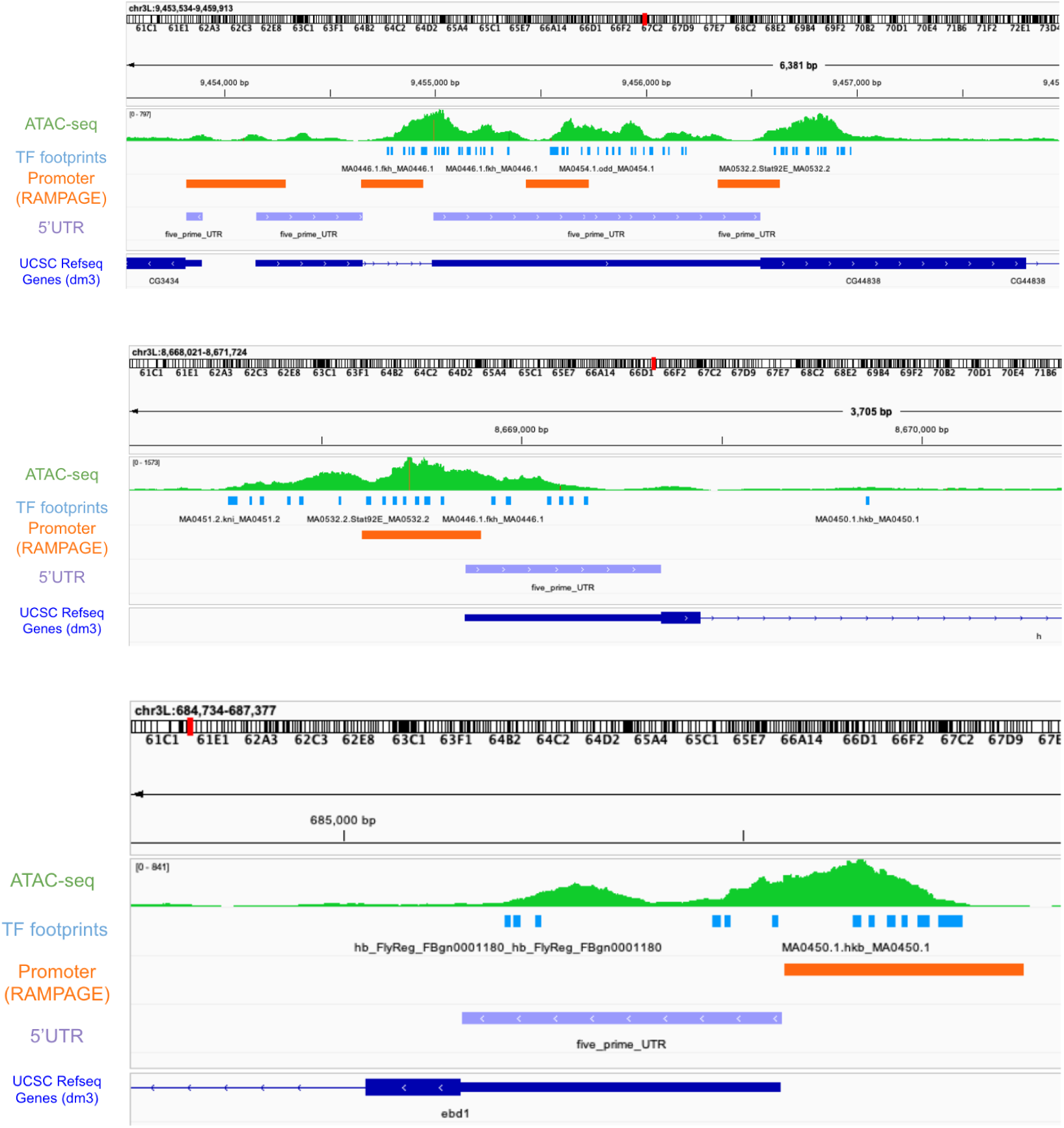
Examples of TF footprints in 5’ UTRs outside of promoters.

## REFERENCES

Andersson R, Sandelin A. 2020. Determinants of enhancer and promoter activities of regulatory elements. Nat Rev Genet. 21(2):71–87. 10.1038/s41576-019-0173-8

Arnold CD et al. 2013. Genome-Wide Quantitative Enhancer Activity Maps Identified by STARR-seq. Science. 339(6123):1074–1077. 10.1126/science.1232542

Arnold CD et al. 2014. Quantitative genome-wide enhancer activity maps for five Drosophila species show functional enhancer conservation and turnover during cis-regulatory evolution. Nature genetics. 46(7):685–692. 10.1038/ng.3009

Batut PJ, Gingeras TR. 2017. Conserved noncoding transcription and core promoter regulatory code in early Drosophila development. eLife. 6:781. 10.7554/elife.29005

Bentsen M et al. 2020. ATAC-seq footprinting unravels kinetics of transcription factor binding during zygotic genome activation. Nat Commun. 11(1):4267. 10.1038/s41467-020-18035-1

Berman BP et al. 2002. Exploiting transcription factor binding site clustering to identify cis-regulatory modules involved in pattern formation in the Drosophila genome. Proc Natl Acad Sci. 99(2):757–762. 10.1073/pnas.231608898

Boyle AP et al. 2008. High-Resolution Mapping and Characterization of Open Chromatin across the Genome. Cell. 132(2):311–322. 10.1016/j.cell.2007.12.014

Bozek M et al. 2019. ATAC-seq reveals regional differences in enhancer accessibility during the establishment of spatial coordinates in the Drosophila blastoderm. Genome Res. 29(5):771–783. 10.1101/gr.242362.118

Buenrostro JD et al. 2013. Transposition of native chromatin for fast and sensitive epigenomic profiling of open chromatin, DNA-binding proteins and nucleosome position. Nature cell biology. 10(12):1213–1218. 10.1038/nmeth.2688

Burz DS, Rivera-Pomar R, Jäckle H, Hanes SD. 1998. Cooperative DNA-binding by Bicoid provides a mechanism for threshold-dependent gene activation in the Drosophila embryo. EMBO J. 17(20):5998–6009. 10.1093/emboj/17.20.5998

Chen J et al. 2023. Prevalent use and evolution of exonic regulatory sequences in the human genome. Nat Sci. 3(2). 10.1002/ntls.20220058

Chen L, Fish AE, Capra JA. 2018. Prediction of gene regulatory enhancers across species reveals evolutionarily conserved sequence properties. PLoS Comput Biol. 14(10):e1006484. 10.1371/journal.pcbi.1006484

Córdova G et al. 2015. SMAD3 and SP1/SP3 Transcription Factors Collaborate to Regulate Connective Tissue Growth Factor Gene Expression in Myoblasts in Response to Transforming Growth Factor β. J Cell Biochem. 116(9):1880–1887. 10.1002/jcb.25143

Creyghton MP et al. 2010. Histone H3K27ac separates active from poised enhancers and predicts developmental state. Proc Natl Acad Sci. 107(50). 10.1073/pnas.1016071107

Gasperini M, Tome JM, Shendure J. 2020. Towards a comprehensive catalogue of validated and target-linked human enhancers. Nat Rev Genet. 21(5):292–310. 10.1038/s41576-019-0209-0

Halfon MS, Zhu Q, Brennan ER, Zhou Y. 2011. Erroneous attribution of relevant transcription factor binding sites despite successful prediction of cis-regulatory modules. BMC Genomics. 12:578. 10.1186/1471-2164-12-578

Hammal F et al. 2021. ReMap 2022: a database of Human, Mouse, Drosophila and Arabidopsis regulatory regions from an integrative analysis of DNA-binding sequencing experiments. Nucleic Acids Res. 50(D1):D316–D325. 10.1093/nar/gkab996

Hardison RC, Taylor J. 2012. Genomic approaches towards finding cis-regulatory modules in animals. Nature Reviews Genetics. 13(7):469–483. 10.1038/nrg3242

Hare EE et al. 2008. Sepsid even-skipped enhancers are functionally conserved in Drosophila despite lack of sequence conservation. PLoS genetics. 4(6):e1000106. 10.1371/journal.pgen.1000106

Hesselberth JR et al. 2009. Global mapping of protein-DNA interactions in vivo by digital genomic footprinting. Nat Methods. 6(4):283–289. 10.1038/nmeth.1313

Jayaram N, Usvyat D, Martin ACR. 2016. Evaluating tools for transcription factor binding site prediction. BMC Bioinform. 17(1):547. 10.1186/s12859-016-1298-9

Johnson DS, Mortazavi A, Myers RM, Wold B. 2007. Genome-Wide Mapping of in Vivo Protein-DNA Interactions. Science. 316(5830):1497–1502. 10.1126/science.1141319

Keränen SVE et al. 2006. Three-dimensional morphology and gene expression in the Drosophila blastoderm at cellular resolution II: dynamics. Genome Biology. 7(12):R124. 10.1186/gb-2006-7-12-r124

Keränen SVE, Villahoz-Baleta A, Bruno AE, Halfon MS. 2022. REDfly: An Integrated Knowledgebase for Insect Regulatory Genomics. Insects. 13(7):618. 10.3390/insects13070618

Koenecke N et al. 2017. Drosophila poised enhancers are generated during tissue patterning with the help of repression. Genome Res. 27(1):64–74. 10.1101/gr.209486.116

Kvon EZ et al. 2014. Genome-scale functional characterization of Drosophila developmental enhancers in vivo. Nature. [published online ahead of print]. 10.1038/nature13395

Lambert SA et al. 2018. The Human Transcription Factors. Cell. 172(4):650–665. 10.1016/j.cell.2018.01.029

Lebrecht D et al. 2005. Bicoid cooperative DNA binding is critical for embryonic patterning in Drosophila. Proceedings of the National Academy of Sciences of the United States of America. 102(37):13176–13181. 10.1073/pnas.0506462102

Lécuyer E et al. 2007. Global Analysis of mRNA Localization Reveals a Prominent Role in Organizing Cellular Architecture and Function. Cell. 131(1):174–187. 10.1016/j.cell.2007.08.003

Li X-Y et al. 2008. Transcription factors bind thousands of active and inactive regions in the Drosophila blastoderm. PLoS Biol. 6(2):e27. 10.1371/journal.pbio.0060027

Li X-Y et al. 2014. Establishment of regions of genomic activity during the Drosophila maternal to zygotic transition. eLife. 3. 10.7554/elife.03737

Li Z et al. 2019. Identification of transcription factor binding sites using ATAC-seq. Genome Biol. 20(1):45. 10.1186/s13059-019-1642-2

Lim LWK, Chung HH, Chong YL, Lee NK. 2018. A survey of recently emerged genome-wide computational enhancer predictor tools. Comput Biol Chem. 74:132–141. 10.1016/j.compbiolchem.2018.03.019

Long HK, Prescott SL, Wysocka J. 2016. Ever-Changing Landscapes: Transcriptional Enhancers in Development and Evolution. Cell. 167(5):1170–1187. 10.1016/j.cell.2016.09.018

Ludwig MZ et al. 2005. Functional evolution of a cis-regulatory module. PLoS Biol. 3(4):e93. 10.1371/journal.pbio.0030093

McGregor AP. 2005. How to get ahead: the origin, evolution and function of bicoid. BioEssays. 27(9):904–913. 10.1002/bies.20285

Moudgil A, Sobti RC, Kaur T. 2023. In-silico identification and comparison of transcription factor binding sites cluster in anterior-posterior patterning genes in Drosophila melanogaster and Tribolium castaneum. PLOS ONE. 18(8):e0290035. 10.1371/journal.pone.0290035

Ohler U, Liao G, Niemann H, Rubin GM. 2002. Computational analysis of core promoters in the Drosophila genome. Genome Biology. 3(12):RESEARCH0087

Öztürk-Çolak A et al. 2024. FlyBase: updates to the Drosophila genes and genomes database. Genetics. 227(1):iyad211. 10.1093/genetics/iyad211

Pfeiffer BD et al. 2008. Tools for neuroanatomy and neurogenetics in Drosophila. Proc Natl Acad Sci. 105(28):9715–9720. 10.1073/pnas.0803697105

Qi Z et al. 2022. Large-scale analysis of Drosophila core promoter function using synthetic promoters. Mol Syst Biol. 18(2):MSB20209816. 10.15252/msb.20209816

Quinlan AR, Hall IM. 2010. BEDTools: a flexible suite of utilities for comparing genomic features. Bioinformatics. 26(6):841–842. 10.1093/bioinformatics/btq033

Rauluseviciute I et al. 2023. JASPAR 2024: 20th anniversary of the open-access database of transcription factor binding profiles. Nucleic Acids Res. 52(D1):D174–D182. 10.1093/nar/gkad1059

Ryczek N, Łyś A, Makałowska I. 2023. The Functional Meaning of 5′UTR in Protein-Coding Genes. Int J Mol Sci. 24(3):2976. 10.3390/ijms24032976

Schulz KN et al. 2015. Zelda is differentially required for chromatin accessibility, transcription factor binding, and gene expression in the early Drosophila embryo. Genome Res. 25(11):1715–1726. 10.1101/gr.192682.115

Shlyueva D, Stampfel G, Stark A. 2014. Transcriptional enhancers: from properties to genome-wide predictions. Nat Rev Genet. 15(4):272–286 http://www.nature.com.ezp-prod1.hul.harvard.edu/nrg/journal/v15/n4/full/nrg3682.html. 10.1038/nrg3682

Smith GD, Ching WH, Cornejo-Páramo P, Wong ES. 2023. Decoding enhancer complexity with machine learning and high-throughput discovery. Genome Biol. 24(1):116. 10.1186/s13059-023-02955-4

Spivakov M. 2014. Spurious transcription factor binding: Non-functional or genetically redundant? BioEssays. 36(8):798–806. 10.1002/bies.201400036

Sun Y et al. 2015. Zelda overcomes the high intrinsic nucleosome barrier at enhancers during Drosophila zygotic genome activation. Genome Res. 25(11):1703–1714. 10.1101/gr.192542.115

Sung M-H, Baek S, Hager GL. 2016. Genome-wide footprinting: ready for prime time? Nat Methods. 13(3):222–228. 10.1038/nmeth.3766

Tomancak P et al. 2002. Systematic determination of patterns of gene expression during Drosophila embryogenesis. Genome Biol. 3(12):research0088.1-88.14. 10.1186/gb-2002-3-12-research0088

Villar D et al. 2015. Enhancer evolution across 20 mammalian species. Cell. 160(3):554–566 http://linkinghub.elsevier.com/retrieve/pii/S0092867415000070. 10.1016/j.cell.2015.01.006

Wunderlich Z, Mirny LA. 2009. Different gene regulation strategies revealed by analysis of binding motifs. Trends in genetics : TIG. 25(10):434–440. 10.1016/j.tig.2009.08.003

Xie X et al. 2020. Smad3 Regulates Neuropilin 2 Transcription by Binding to its 5′ Untranslated Region. J Am Hear Assoc. 9(8):e015487. 10.1161/jaha.119.015487

Yan F, Powell DR, Curtis DJ, Wong NC. 2020. From reads to insight: a hitchhiker’s guide to ATAC-seq data analysis. Genome Biol. 21(1):22. 10.1186/s13059-020-1929-3

Yang T-H, Yang Y-C, Tu K-C. 2022. regCNN: identifying Drosophila genome-wide cis-regulatory modules via integrating the local patterns in epigenetic marks and transcription factor binding motifs. Comput Struct Biotechnol J. 20:296–308. 10.1016/j.csbj.2021.12.015

Zhu LJ et al. 2011. FlyFactorSurvey: a database of Drosophila transcription factor binding specificities determined using the bacterial one-hybrid system. Nucleic Acids Res. 39(suppl_1):D111–D117. 10.1093/nar/gkq858

